# Early diverging fungus *Mucor circinelloides* lacks centromeric histone CENP-A and displays a mosaic of point and regional centromeres

**DOI:** 10.1101/706580

**Authors:** María Isabel Navarro-Mendoza, Carlos Pérez-Arques, Shweta Panchal, Francisco E. Nicolás, Stephen J. Mondo, Promit Ganguly, Jasmyn Pangilinan, Igor V. Grigoriev, Joseph Heitman, Kaustuv Sanyal, Victoriano Garre

## Abstract

Centromeres are rapidly evolving across eukaryotes, despite performing a conserved function to ensure high fidelity chromosome segregation. CENP-A chromatin is a hallmark of a functional centromere in most organisms. Due to its critical role in kinetochore architecture, the loss of CENP-A is tolerated in only a few organisms, many of which possess holocentric chromosomes. Here, we characterize the consequence of the loss of CENP-A in the fungal kingdom. *Mucor circinelloides*, an opportunistic human pathogen, lacks CENP-A along with the evolutionarily conserved CENP-C, but assembles a monocentric chromosome with a localized kinetochore complex throughout the cell cycle. Mis12 and Dsn1, two conserved kinetochore proteins were found to bind nine short overlapping regions, each comprising an ∼200-bp AT-rich sequence followed by a centromere-specific conserved motif that echoes the structure of budding yeast point centromeres. Resembling fungal regional centromeres, these core centromere regions are embedded in large genomic expanses devoid of genes yet marked by Grem-LINE1s, a novel retrotransposable element silenced by the Dicer-dependent RNAi pathway. Our results suggest that these hybrid features of point and regional centromeres arose from the absence of CENP-A, thus defining novel mosaic centromeres in this early-diverging fungus.

## Introduction

Accurate chromosome segregation is crucial to maintain genome integrity during cell division. The timely attachment of microtubules to centromere DNA is essential to achieve proper chromosome segregation. This is accomplished by a specialized multilayered protein complex, the kinetochore which links microtubules to centromere DNA. This protein bridge is divided into two layers – the inner and outer kinetochore. The fundamental inner kinetochore protein is the histone H3 variant CENP-A. It binds directly to centromere DNA and lays the foundation to recruit other essential proteins of the kinetochore complex, playing a fundamental role in centromere structure and function, and hence, precise chromosome segregation^1,2^. CENP-A is also found at all identified neocentromeres^3^ and at the active centromeres of dicentric chromosomes^4^, acting as an epigenetic determinant of centromeric identity.

Despite its conserved function, the centromere is one of the most rapidly evolving regions of the genome^5^. This so-called “centromere paradox” has led to centromeres of diverse sizes and content. The first centromeres identified in *Saccharomyces cerevisiae* were found to be point centromeres - small regions of ∼120 bp defined by specific DNA sequences^6,7^. In contrast to point centromeres described in only a few budding yeasts of the phylum Ascomycota, most other fungi and metazoans have regional centromeres that are larger, ranging from a few kilobases to several megabases^8^. Regional centromeres are often interspersed with repetitive sequences and are mostly defined by epigenetic factors rather than DNA sequence per se^9^. While the centromeres described thus far are confined to one region of the DNA, there are organisms in which their entire chromosomes are loaded with kinetochores and display a parallel separation of the two chromatids. Such chromosomes possess holocentromeres and have been found in several clades of insects^10^ and nematodes like *Caenorhabditis elegans*^11^. Despite this remarkable variation in the types of centromeres, most organisms have their centromere loci defined by the binding of CENP-A. However, there are a few exceptions, including some insect lineages and kinetoplastids, which have independently lost CENP-A. The kinetoplastid *Trypanosoma brucei* has evolved a unique set of proteins to perform the role of the kinetochore complex^12^, whereas all insects that have recurrently lost CENP-A have transitioned from monocentricity to holocentricity^10^.

Neither holocentricity nor lack of CENP-A has ever been reported in the fungal kingdom until a recent study showed that this protein could be absent in the Mucoromycotina subphylum^13^, which belongs to the early diverging fungi. Kinetochore structure and centromere function are well-studied in species of the Dikarya, whereas centromere identity as well as the kinetochore architecture remain largely unexplored in basal fungi because these are challenging to manipulate genetically. *Mucor circinelloides*, a fungus of the subphylum Mucoromycotina, is an exception and molecular tools are available to modify its genome^14^. *M. circinelloides*, as with other Mucorales, causes an infectious disease known as mucormycosis, which have been associated with high mortality rates due to rhino-orbital-cerebral, pulmonary, or cutaneous infections^15^. Despite its medical significance and research efforts to identify new antifungal targets, the genome biology and cell cycle of this fungus remain poorly understood. Using evolutionarily conserved kinetochore proteins as tools, we investigated the dynamics of the kinetochore complex during nuclear division and unveiled the unique characteristics of the centromeres of *M. circinelloides*. Studying distinct and divergent factors involved in the cellcycle of this organism will open avenues for the development of species-specific antifungal drugs.

## Results

### Last Mucoralean common ancestor lost CENP-A and CENP-C, and the descendant Mucoromycotina lack centromere-specific histone variants

A recent study of kinetochore evolution in eukaryotes reported the striking absence of two well conserved inner kinetochore proteins, CENP-A and CENP-C, in two species of the Mucoromycotina^13^. To ascertain the presence or loss of well-studied kinetochore proteins across the entire subphylum, we analyzed the curated proteomes of 75 fungal species, including 55 species of Mucoromycotina and 20 species from other fungal clades (Supplementary Table 1). The kinetochore protein sequences from *Mus musculus, Ustilago maydis, S. cerevisiae*, and *Schizosaccharomyces pombe* served as queries to conduct iterative Blast and Hmmer searches against these proteomes. These sequences were confirmed as putative orthologs by the presence of conserved Pfam domains or by obtaining a matching first hit in a reciprocal Blastp search against the initial four species: mouse, *U. maydis*, budding, and fission yeasts (Supplementary Table 1, Supplementary Data 1).

To confirm the presence or absence of centromere-specific histone H3 CENP-A in each fungal species, 289 histone H3 orthologs were analyzed, identifying 27 putative CENP-A proteins (Supplementary Fig. 1, Supplementary Table 1, Supplementary Data 1). These proteins share common features with well-known CENP-A proteins^16^, such as an N-terminal tail that varies significantly in length and sequence compared to the canonical H3 counterparts (Supplementary Fig. 1b), the absence of a conserved glutamine residue in the first α-helix (69Q) and a phenylalanine residue (84F) (Supplementary Fig. 1c), a longer and divergent first loop (Supplementary Fig. 1c), and a non-conserved C-terminal tail (Supplementary Fig. 1d). In addition, the <u>h</u>istone <u>f</u>old <u>d</u>omain (HFD) sequences of the putative CENP-A proteins have diverged rapidly from the conserved canonical H3 sequences^16^ (Supplementary Fig. 1e). Nine additional H3 orthologs also share several CENP-A common features but were regarded as rare histones because they lack these conserved substitutions. The analysis revealed that CENP-A was absent in most of the Mucoromycotina clade, specifically in the orders Umbelopsidales and Mucorales; nevertheless, we identified putative CENP-A proteins in all Endogonalean species (Fig. 1). Similarly, CENP-C is present in the Endogonales and Umbelopsidales but not in the Mucorales (Fig. 1). These findings suggest that the last Mucoromycotina common ancestor possessed CENP-A, which was lost after the Endogonales diverged from the rest of the clade. In the absence of CENP-A, CENP-C was presumably lost soon after. In addition, CENP-Q is absent in all Mucoralean species, and CENP-U is lost in the whole subphylum. These proteins have been recurrently lost across the fungal kingdom, as evidenced by their absence in species of all phyla of early-diverging fungi and most species in Basidiomycota (Fig. 1). The absence of orthologs that were present in closely-related species should be taken with caution because this could be a result of the incomplete state of their genome assemblies and annotation.

**Fig 1.**
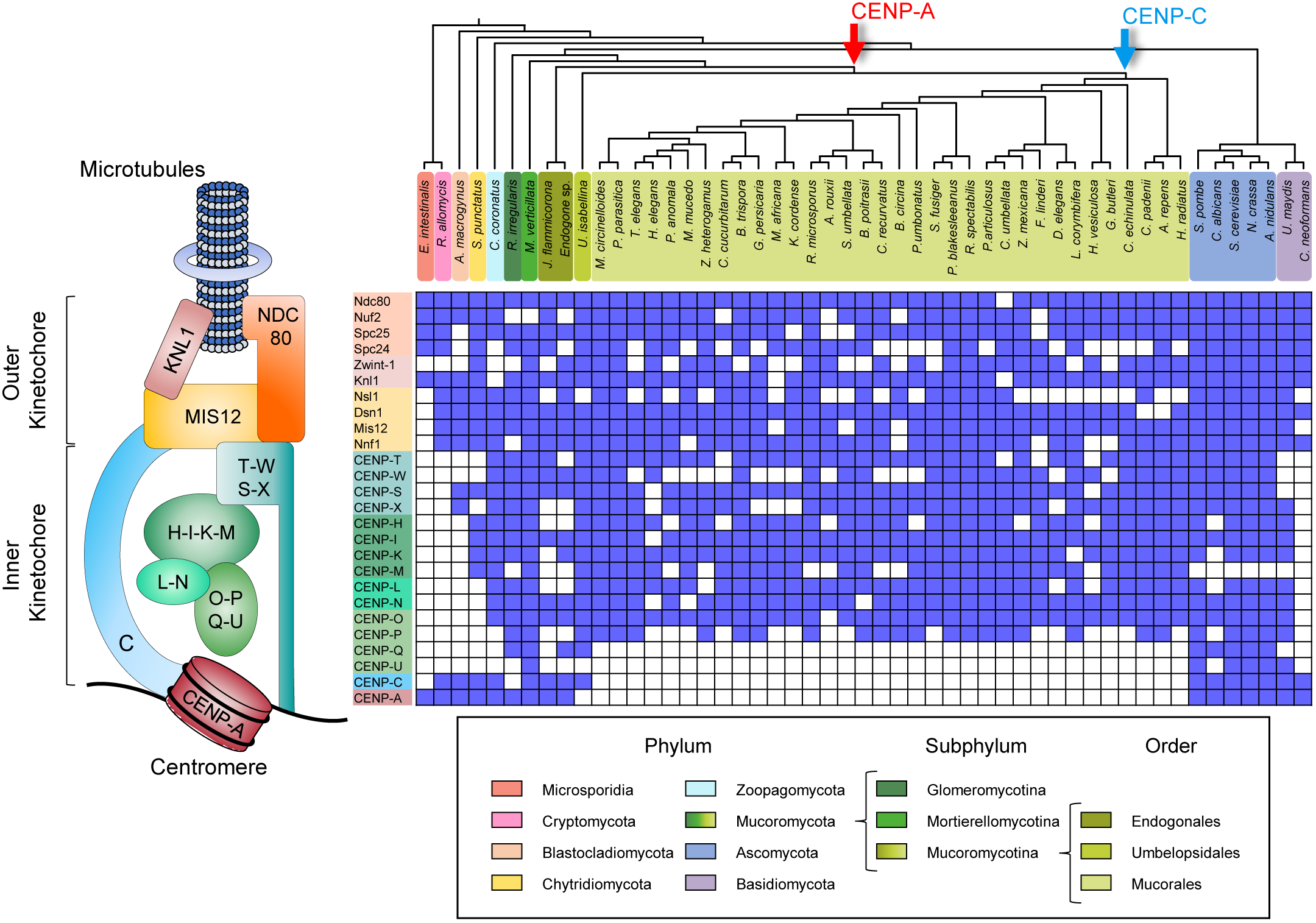
Distribution of the kinetochore complex across the Mucoromycotina subphylum. Concise kinetochore schematic showing the most conserved protein complexes in eukaryotes is shown on the left corresponding to the kinetochore proteins analyzed in the matrix on the right. The matrix displays the presence or absence of 26 kinetochore proteins across a cladogram of 51 fungal lineages (top) to show the relationships among them. Every fungal phylum is represented and color-coded, differentiating the Mucoromycota into the Glomeromycotina, Mortierellomycotina, and Mucoromycotina subphyla to provide a special emphasis on the latter. Arrows at divergence events in the cladogram mark the hypothetical loss of proteins CENP-A (red) and CENP-C (blue) in these clades.

Despite these relevant losses, the remaining kinetochore proteins were well conserved among most species of Mucoromycotina, especially in *M. circinelloides*. Therefore, we hypothesized that another histone protein could have taken the role of CENP-A as a part of centromeric nucleosome for kinetochore assembly. We searched the *M. circinelloides* genome sequence for histone protein-coding genes and examined their sequences seeking distinctive features of histone variants. In contrast to canonical mammalian histones H3.1 and H3.2, fungal cell cycle-dependent histones may have intronic sequences and polyadenylation [poly(A)] signals^17,18^. Thus, the analysis of possible variants focused on the presence of substitutions in relevant amino acid residues targeted by post-translational modifications^19^. The *M. circinelloides* genome contains four histone H3- and three histone H4-coding genes, namedm *<u>h</u>istone <u>H</u> three* (*hht1-4*) and *<u>h</u>istone <u>H</u> four* (*hhf1-3*), respectively (Supplementary Fig. 2a), and all of them feature a poly(A) signal. Three of the four histone H3-coding genes are intronless and their protein sequence identical, except the intron-containing gene *hht4* (Supplementary Fig. 2a). Also, the amino acid residues at positions 32, 43, and 97 in the Hht4 protein sequence differ compared to the conserved H3 sequence (Supplementary Fig. 2b). Similarly, histone H4-coding genes *hhf2* and *hhf3* lack introns and encode identical proteins, whereas *hhf1* has intronic sequences (Supplementary Fig. 2a) and its product has a longer N-terminal tail (Supplementary Fig. 2c).

**Fig 2.**
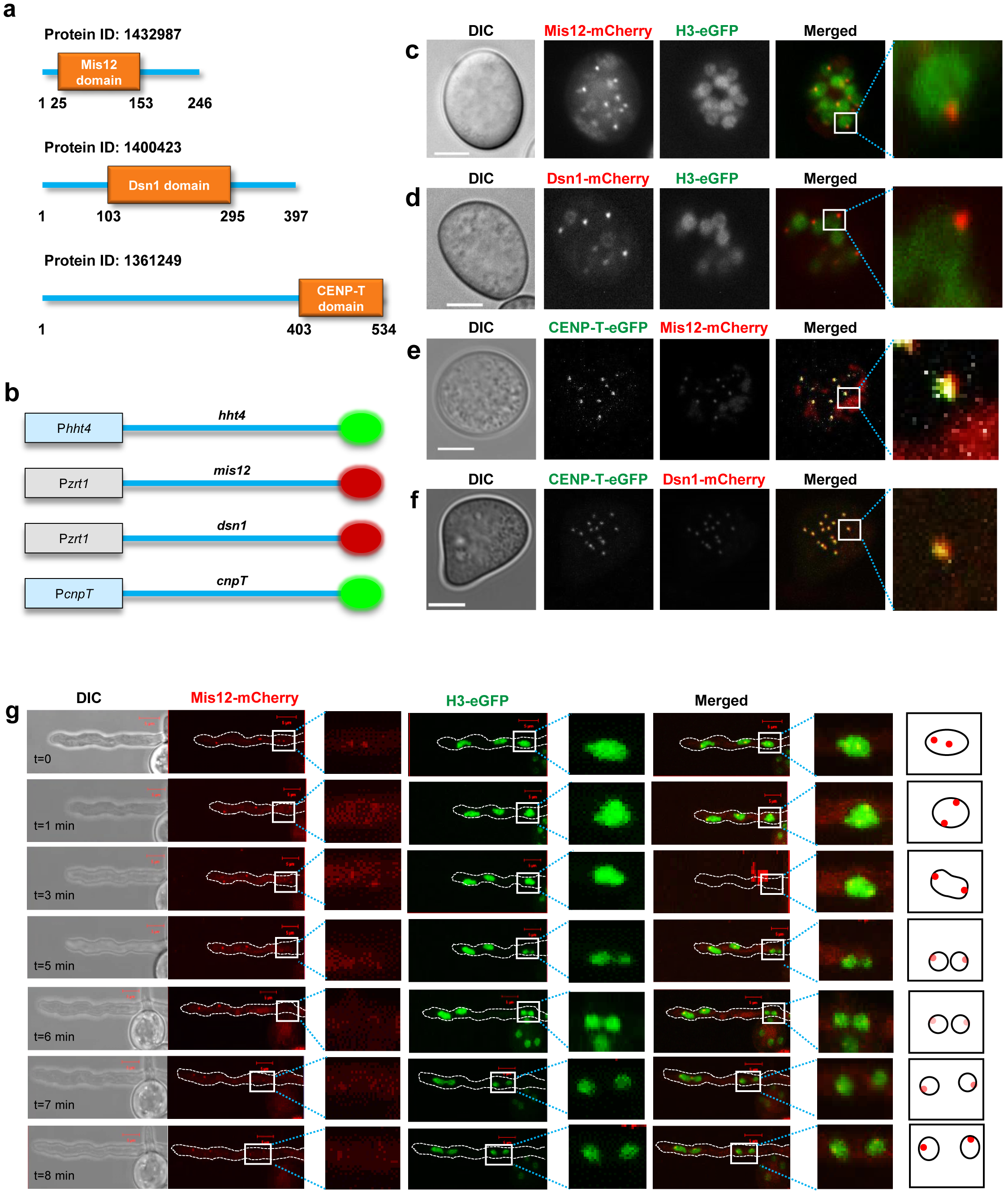
Inner and outer kinetochore proteins colocalize in *M. circinelloides* nuclei in the absence of CENP-A and CENP-C. **(a)** Graphical representation of Mis12, Dsn1, and CENP-T sequence features showing individual Pfam domains PF05859, PF08202, PF15511 respectively identified in *M. circinelloides*. **(b)** Schematic to represent C-terminal tagging of histone H3 and kinetochore proteins with mCherry (red circles), eGFP (green circles). **(c-f)** Confocal microscope imaging showing the cellular localization of well-conserved mCherry-tagged outer kinetochore proteins Mis12 **(c, e)** and Dsn1 **(d, f)**, eGFP-tagged histone H3 **(c, d)**, and eGFP-tagged inner kinetochore CENP-T **(e, f)** in pre-germinated spores of *M. circinelloides* strains expressing fluorescent fusion proteins. A calibrated scale (white bar) is provided for size comparison (5 µm). **(f)** Time-lapsed confocal image displaying the cellular localization of mCherry-tagged Mis12 and eGFP-tagged histone H3 in a germinative tube sprouting from a spore. A discontinuous white perimeter outlines the germinative tube in the fluorescence images, and a yellow circle indicates a nuclear division event. Cartoon for the zoomed image represents the localization and signal intensity of Mis12-mCherry in the nucleus.

To test if the minor differences observed in Hht4 and Hhf1 could have resulted in a centromere-specific binding protein, the cellular localization was analyzed and compared to their canonical histone counterpart Hhf3. We constructed alleles that contained each histone-coding gene fused in-frame at the C-terminus with the enhanced green fluorescent protein (eGFP) flanked by each of the histone gene native promoter and terminator sequences, and a selectable marker (Supplementary Fig. 3). Homokaryotic strains expressing fluorescent histones Hht4, Hhf1, or Hhf3 were obtained by transforming a wild-type strain with their respective eGFP-tagged alleles. Integration by homologous recombination at the corresponding native loci was confirmed by PCR (Supplementary Fig. 3, Supplementary Tables 2,3).

**Fig 3.**
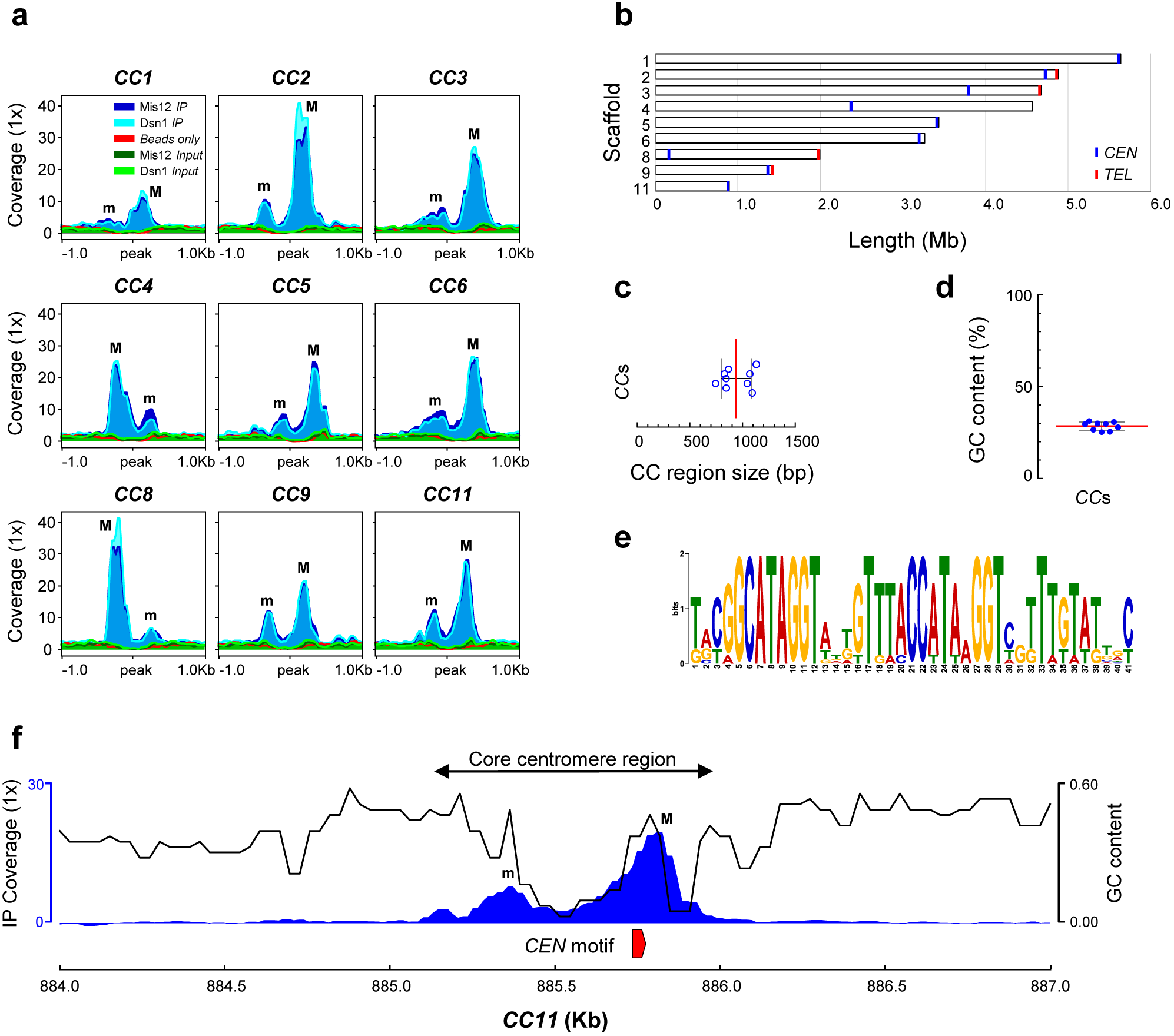
The nine centromeres of *M. circinelloides* are short, AT-rich, and harbor a conserved DNA motif. **(a)** Putative kinetochore-binding regions showing color-coded enrichment of immunoprecipitated DNA (*IP* DNA) from Mis12 and Dsn1 mCherry-tagged strains compared to their corresponding input (*Input* DNA) and binding controls (*Beads only*). 1 kb flanking sequences from the center of the enrichment peak are shown. Major (M) and minor (m) kinetochore-binding regions are indicated. **(b)** Scaled ideogram of the nine scaffolds (white) of *M. circinelloides* genome assembly that contain a kinetochore-binding region (blue), showing the telomeric repeats (red). **(c)** Size distribution of the nine core centromere (CC) regions. **(d)** GC content across the nine core centromere (CC) sequences. The median (red line) and standard deviation (black lines) are shown in **c** and **d. (e)** 41 bp centromere-specific DNA motif conserved in all nine CC regions and absent in the remainder of the genome. **(f)** A genomic view of *CEN11* illustrating all nine core centromere regions. Kinetochore-binding region enrichment (IP coverage) as the average of both immunoprecipitation signals minus *Input* and *beads only* controls (left axis, blue), GC content across the region (right axis, black), and the centromere-specific DNA motif described in **e** (red) are shown.

Asexual spores from these strains were observed with a confocal microscope under two stages of germination: ungerminated and 4-h pre-germinated spores, that had started swelling and forming germ tubes. In both conditions, all fluorescent histone fusions were localized encompassing the entire nucleus instead of forming kinetochore-like nuclear clusters (Supplementary Fig. 2d). Most species of the Mucoromycotina are coenocytic fungi and have multinucleated spores, as does the *M. circinelloides* wild-type strain used in this study which has large spores with a high number of nuclei^20^. We established that nuclear division in *M. circinelloides* is asynchronous by observing the nuclear distribution of Hht4-eGFP protein in pre-germinated spores during a time-lapse imaging session (Supplementary video 1). Altogether, these findings indicate that either none of these histone variants functions like CENP-A in kinetochore assembly or *Mucor* centromeres are holocentric in nature. We could test these two possibilities by studying other kinetochore proteins that are evolutionarily conserved and easily identifiable in *M. circinelloides*.

### Mis12 and CENP-T complexes show discrete nuclear localization

Analyzing the nuclear localization and assembly dynamics of the kinetochore complex of *M. circinelloides* could offer insights into its architecture. In the absence of CENP-A or another similar histone H3 variant, and CENP-C, that could anchor the kinetochore formation, we focused our attention on the evolutionarily conserved and obvious homologs of inner kinetochore proteins that are expected to be closer to the centromere DNA in this organism (Fig. 2a). We designed fluorescent kinetochore fusion proteins of Mis12, Dsn1, and CENP-T because these are well-conserved in almost all species of the Mucoromycotina. First, we constructed alleles to tag the Mis12 and Dsn1 proteins with the red fluorescent protein mCherry, both at their C-termini and N-termini, aiming to integrate the alleles in their endogenous loci following a similar strategy as for the epitope tagging of the histone genes described above. Homologous recombination in the desired transformants was confirmed by PCR, and six mutant colonies from the four tagged alleles were checked for fluorescent localization. Unfortunately, fluorescent signals were not detected in any of the four different tagged strains possibly due to low levels of expression.

As an alternative approach, the Mis12 and Dsn1 proteins were tagged with mCherry at both ends and overexpressed from a strong promoter (P*zrt1*) in *M. circinelloides* (Fig. 2b, Supplementary Fig. 4). The alleles were designed to target integration into the *carRP* locus, a gene encoding an enzyme involved in carotenogenesis. Thus, the mutant strains obtained should lack colored-carotenes and display an albino phenotype^14^. After confirming the integration of the alleles in the *carRP* locus by PCR (Supplementary Fig. 4a, b, Supplementary Table 2), one strain harboring each construct was used as a parental strain to integrate the allele for H3-eGFP expression at its own locus as previously described, allowing monitoring colocalization of the kinetochore proteins within the nucleus. Once the double integration was confirmed, two independent strains of each type were selected for fluorescent localization (Supplementary Table 3).

**Fig 4.**
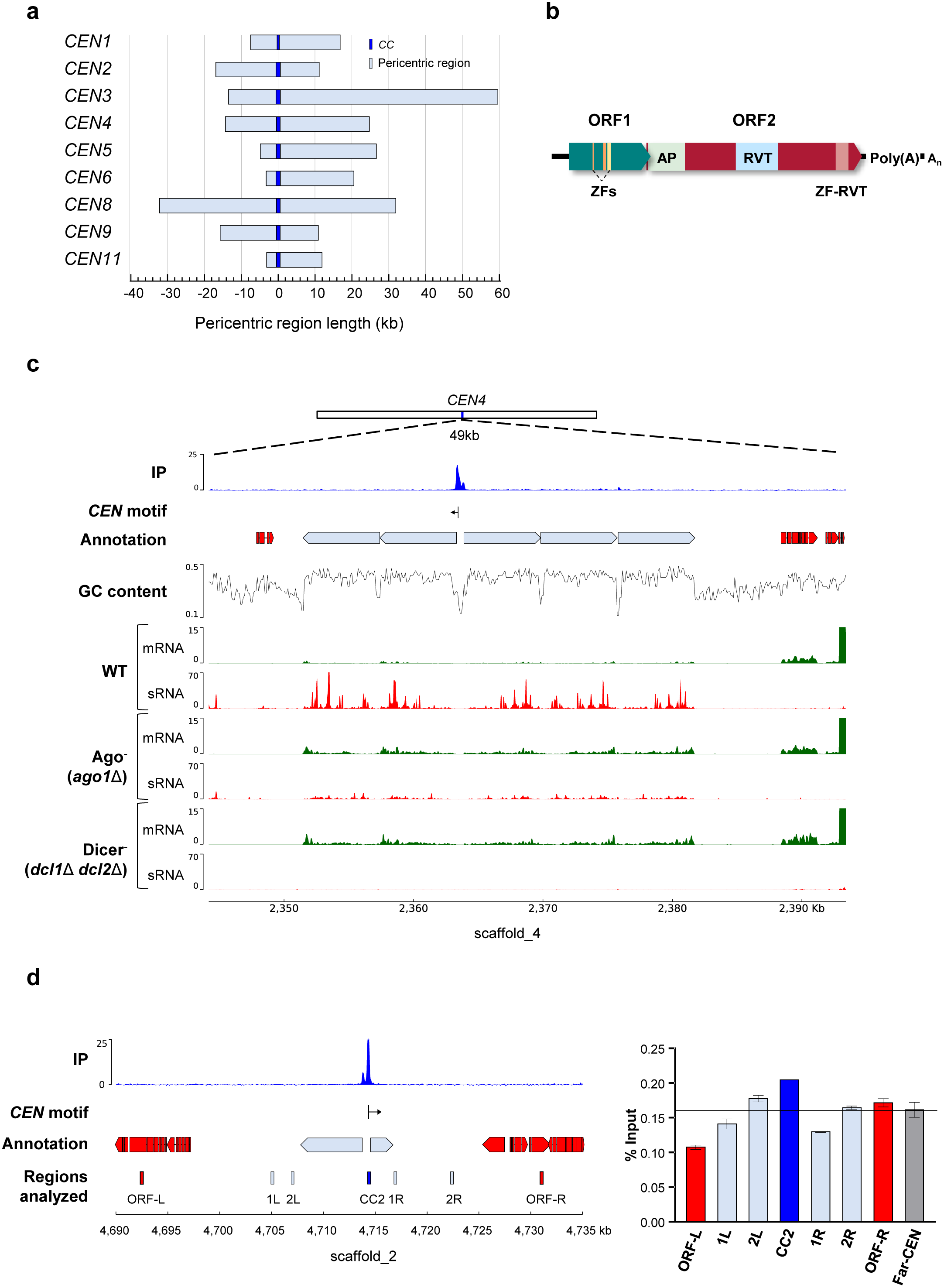
*M. circinelloides* core centromeres are present in large ORF-free pericentric regions having retrotransposable elements regulated by the Dicer-dependent RNAi pathway and bind H3 nucleosomes. **(a)** Genomic sequences lacking annotated genes (light blue) that flank the *CC* regions (dark blue) served as the reference points to calculate the length both upstream and downstream. **(b)** Schematic of the Grem-LINE1 interspersed in the pericentromeric regions. <u>O</u>pen <u>R</u>eading <u>F</u>rames (ORF) and protein domains predicted from their coding sequences are shown as colored boxes [Zf, zinc finger (PF00098 and PF16588); AP, AP endonuclease (PTHR22748); RVT, reverse transcriptase (PF00078); and zf-RVT, zinc-binding in reverse transcriptase (PF13966)]. **(c)** *CEN4* is exemplified by low GC content, lack of genes and transcripts, which are shown in different tracks displaying the kinetochore-binding region enrichment (IP, an average of both *IP* DNA signals minus *Input* and *beads only* controls), annotation of genes (red blocks) and transposable elements (light blue blocks), *CEN*-specific DNA motif position (vertical line) and direction (arrow), GC content, and transcriptomic data of mRNA (green) and sRNAs (red) in *M. circinelloides* wild-type, *ago1*, and double *dcl1 dcl2* deletion mutant strains after 48 h of growth in rich media. **(d)** Genomic location of *CEN2* showing the regions studied for histone H3 occupancy as labelled and colored rectangles. ChIP assays were performed using polyclonal antibodies against histone H3. Primers were designed for *CC2* region, pericentric regions (1L, 2L, 1R, 2R), flanking ORFs (ORF-L, ORF-R) and a far-*CEN* control ORF ∼2 Mb away from *CEN2* (Far-*CEN*). *IP* samples were analyzed by real-time PCR using these primers. The *y*-axis denotes the qPCR value as a percentage of the total chromatin input with standard error mean (SEM), from each region tested. The experiment was repeated three times with similar results.

Mis12 and Dsn1 displayed a clear localization signal forming small fluorescent puncta within the nucleus marked by histone H3 (Fig 2c, d respectively). The microscopy screening confirmed that the expression of both N-terminal and C-terminal tagged proteins was similar. The Mis12 and Dsn1 localization patterns revealed a single cluster of kinetochores as observed in many fungal species. This result strongly suggests that *M. circinelloides* has monocentric chromosomes with a localized kinetochore even in the absence of CENP-A.

Next, another evolutionarily conserved inner kinetochore protein CENP-T, which is known to interact with DNA, was tagged with eGFP at the C-terminus following the same procedure used for the histone tagging and integrated by homologous recombination at the endogenous locus in strains expressing the Mis12-mCherry and Dsn1-mCherry fusion proteins. Integration of the alleles at the desired loci was confirmed by PCR in both parental backgrounds (Supplementary Fig. 4c), and the localization of the kinetochore proteins was examined by analyzing their fluorescent signal in pre-germinated spores. CENP-T colocalized with Mis12 and Dsn1 in each nucleus (Fig 2e, f, respectively), indicating that all three proteins assemble together in the kinetochore ensemble. Time-lapse imaging during spore germination of the double-tagged strains allowed the analysis of nuclear division and kinetochore structure during the cell cycle in live cells. The Mis12 and Dsn1 proteins clustered at the nuclear periphery during division (Fig 2g), a typical feature of fungal kinetochore proteins, suggesting all these three proteins are constitutively present at the kinetochore during the cell cycle.

### *M. circinelloides* exhibits small mosaic centromeres associated with RNAi-suppressed L1-like retrotransposons

The centromere regions of early-diverging fungi have never been described, possibly because of their genetic intractability. The generation of kinetochore-tagged strains of *M. circinelloides* offered the opportunity to identify the nature of centromeres in an organism that lost CENP-A and CENP-C. To map *M. circinelloides* centromeres, <u>ch</u>romatin immuno<u>p</u>recipitation followed by next generation <u>seq</u>uencing (ChIP-seq) was performed with Mis12- and Dsn1-mCherry proteins. Illumina sequencing was conducted in two independently immunoprecipitated (*IP* DNA) samples for each kinetochore protein, one for Mis12-mCherry and the other for Dsn1-mCherry, using RFP-Trap MA beads (Chromotek) for mCherry immunoprecipitation. Input control samples consisting of total sheared DNA (*Input* DNA) were also sequenced for each tagged strain, as well as mock binding control (*beads only* DNA) samples with MA beads without antibody. Raw sequencing data were processed, and reads were aligned to a newly generated PacBio genome assembly of *Mucor circinelloides* f. *lusitanicus* MU402 strain (hereafter the *M. circinelloides* genome), the parental strain of the kinetochore protein-tagged strains.

Both Mis12 and Dsn1 *IP* DNA sequence reads were compared to the corresponding control *Input* DNA reads to identify statistically significant kinetochore protein-enriched genomic loci in both *IP* DNA samples (FDR ≤ 5×10^−5^, fold enrichment ≥ 1.6; Supplementary Table 4). The result of this analysis was visualized in a genome browser to ensure that the enrichment was robust and absent in either the *Input* or *beads only* DNA controls (Supplementary Fig. 5). By this approach, we detected nine significantly enriched peaks that overlapped in both Mis12 and Dsn1 *IP* DNA samples, and thus define core centromeres (CCs) (Fig 3a). Each of these core *CEN* sequences was located in a different scaffold of the genome assembly (Fig. 3b) and was designated by their scaffold number. The identification of five repeated regions matching the telomeric sequence (5’-TTAGGG-3’) at the 3’-end of scaffolds 2, 3, 8, 9, and 10, suggested that *CC2* and *CC9* display a subtelomeric localization (Fig. 3b). Because the length of the overlapping peaks was minimally different in the two kinetochore *IP* DNA samples, DNA sequence enriched with at least one of the kinetochore proteins was assumed to be the length of kinetochore binding region on each chromosome. In addition, contiguous enriched regions were added to each *CC*, defining the core centromere regions (Supplementary Table 4).

**Fig 5.**
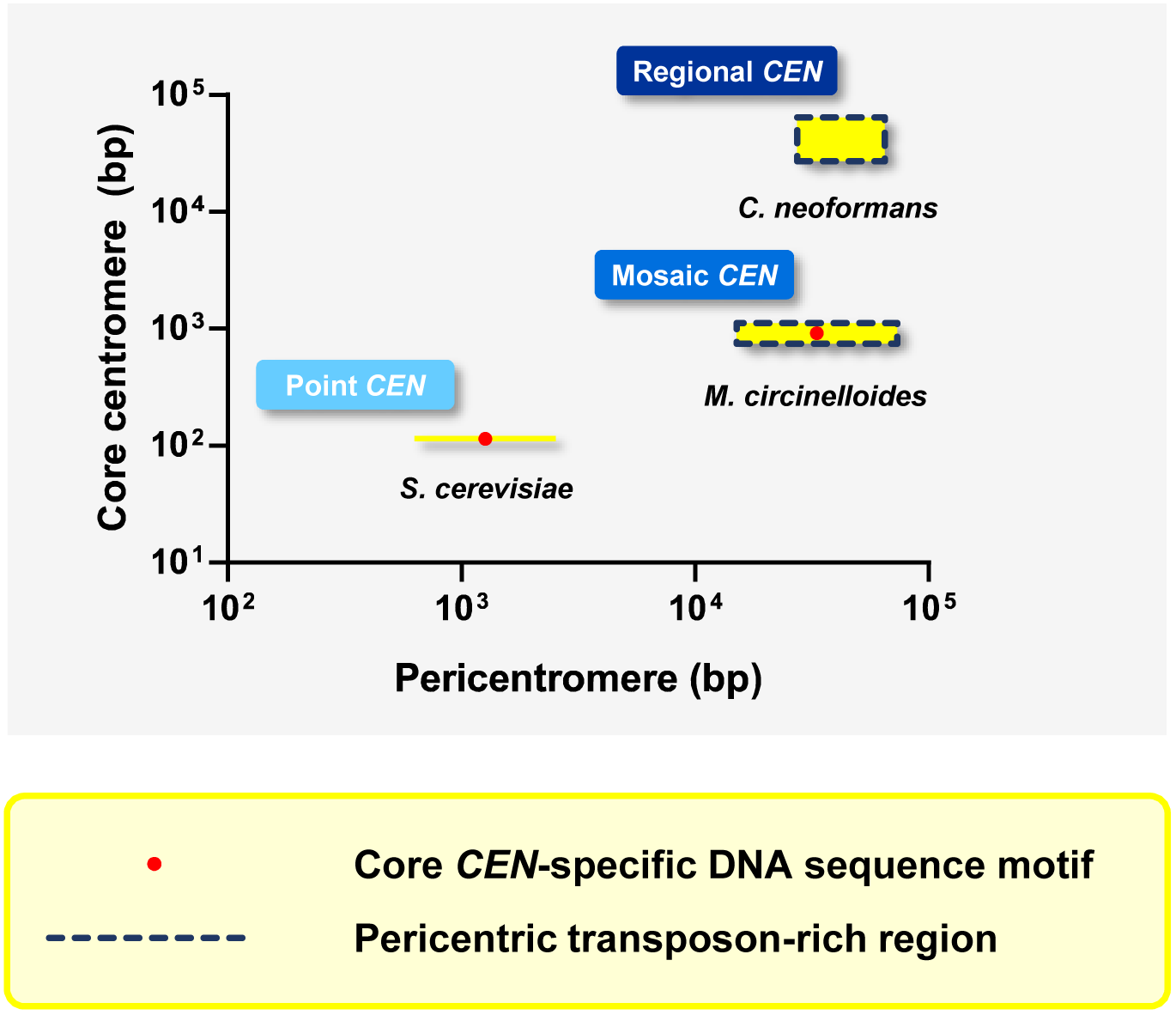
*M. circinelloides* possesses a mosaic of point and regional centromeres. A comparative analysis of centromere features of *S. cerevisiae* (ascomycete), *M. circinelloides* (mucoromycete), and *Cryptococcus neoformans* (basiodiomycete). *M. circinelloides* centromeres are a mosaic with features of point centromeres like shorter kinetochore-bound region (core centromere) and a conserved motif (red dots), combined with features of regional centromeres like large ORF-free pericentromeres dotted with transposable elements that are suppressed by the RNAi machinery.

The average length of core centromere regions of *M. circinelloides* is 941 bp (Fig. 3c). These unusually short regions are consistently AT-rich (71% average) (Fig. 3d), resembling point centromeres of *S. cerevisiae* and its related budding yeast species. Further analysis revealed a 41-bp DNA sequence motif conserved in all nine kinetochore protein-bound core centromere regions (Fig. 3e). An extensive search of the presence of this motif across the whole genome revealed that the 41-bp motif is centromere-specific in *M. circinelloides*. Moreover, a closer inspection of the core centromere regions revealed a conserved pattern. Each core centromere is composed of a minor peak enriched of kinetochore proteins spaced by a highly AT-rich stretch spanning the major binding peak of kinetochore proteins that starts with the 41-bp conserved motif (Fig. 3f).

Surrounding the core centromere regions, both upstream and downstream, are stretches of genomic sequences completely devoid of genes. These pericentric regions are considerably larger than the core and vary in length, ranging from 15 to 75 kb (Fig. 4a). These pericentric regions contain a variable number of large repeats oriented away from the core centromere regions (Supplementary Fig. 6). These repeats cluster in nine groups according to their similarity (≥ 95% identity and ≥ 70% coverage across their entire sequence). Clusters 1, 2 and 3 contain all of the full-length elements, which were termed autonomous elements, while the remaining clusters are composed of incomplete repeats or remnants (Supplementary Fig. 7a). A pairwise comparison of all of the repeats revealed a high similarity in their aligned sequences (≥ 90% identity), though the truncated sequences of remnants showed higher differences accounting for the gaps in the alignment.

Each autonomous repeat comprises two sequential <u>o</u>pen <u>r</u>eading frame<u>s</u> (ORFs) with an overlapping codon. The first ORF encodes a product containing several zinc finger domains, and the second codes for a protein with AP endonuclease and reverse transcriptase domains (Fig 4b, Supplementary Table 5). Based on their high similarity, their ORF domain architecture, the presence of a poly(A) signal at their 3’-end, and the absence of long terminal repeats (LTR) at either end, they were classified as a single non-LTR retrotransposon. Phylogenetic profiling revealed that these retrotransposons belong to the LINE1 clade (Supplementary Fig. 7b) and are closely related to other LINE1-like retrotransposons of Mucoromycotina species, so they were termed <u>G</u>enomic <u>R</u>etrotransposable <u>E</u>lement of <u>M</u>ucoromycotina (Grem) LINE1-like. A search of the *M. circinelloides* genome revealed that the Grem-LINE1s are found specifically in the pericentromeric regions; or at the non-telomeric ends of three small scaffolds, oriented away from the end (Supplementary Table 5). Four major scaffolds end abruptly in a putative centromere region, suggesting that they may be linked to these minor scaffolds that contain the Grem elements at one end. Broadening the search across 55 genomes of species of Mucoromycotina identified the Grem elements in most Mucorales and Umbelopsidales (≥ 30% identity and ≥ 50% coverage), but they could not be found in any of the Endogonalean genomes, indicating an inverse correlation between the presence of Grem-LINE1s and CENP-A (Fig. 1 and Supplementary Table 6).

RNAi silences expression and prevents transposition of retrotransposable elements in several fungal species featuring functional RNAi^21,22^. *M. circinelloides* possess an elaborate RNAi mechanism that protects the genome against invading elements, that could prevent the movement of retrotransposable elements. The two Dicer enzymes, particularly Dcl2, and one of the three Argonaute proteins (Ago1), play a pivotal role in this protective RNAi-related pathway^23^. To analyze the transcriptional landscape of the pericentric regions, previously generated and publicly available transcriptomic raw data for mRNA and small RNAs were reprocessed and aligned to the *M. circinelloides* genome assembly. The pericentric regions were almost devoid of transcription in the wild-type strain, although the retrotransposons were modestly expressed. Indeed, a high number of sRNAs corresponding to the retrotransposons were detected (Fig. 4c, Supplementary Fig. 6), indicating an inverse correlation between mRNA and sRNA levels. This inverse correlation was confirmed by analyzing the transcriptomic profile of mutant strains lacking essential components of the RNAi machinery, Ago1 or Dcl1/Dcl2. In these RNAi-deficient mutants, the pattern was reversed; high mRNA levels from the retrotransposons were observed, while the production of sRNAs originating from those regions was considerably lower than in the wild-type strain. These findings suggest that the retrotransposons contained in the pericentric regions of *M. circinelloides* are silenced by the Dicer-dependent RNAi pathway.

In most organisms, deposition of CENP-A at the centromere-specific nucleosomes is correlated to depletion of histone H3^24–26^. Because CENP-A is absent in *M. circinelloides*, we quantified the presence of histone H3 at the centromere-specific nucleosomes by performing ChIP with anti-histone H3 antibodies and a qPCR analysis. The occupancy of histone H3 was examined by qPCR in four regions across the scaffold_2: the spanning kinetochore protein-binding region (core centromere, *CC2*), the ORF-free pericentromeric region, the centromere-flanking ORFs and the control region far-CEN ORF ∼2 Mb away from the centromere (Fig. 4d). There are no significant differences in histone H3 occupancy across the analyzed regions, indicating that H3 is not depleted at either the kinetochore protein binding region or the pericentric region compared to the flanking ORFs and the control far-CEN ORF (Fig. 4d).

## Discussion

Mucoralean species are the causative agents of mucormycosis, an emerging fungal infection with extremely poor clinical outcomes, probably as a consequence of their innate resistance to all current antifungal drugs^27,28^ and their ability to evade host innate immunity^29,30^. In spite of their clinical relevance, limited information is available on essential biological processes in these fungal pathogens, especially nuclear division and cell cycle. In this study, we identified major kinetochore components in *M. circinelloides*, and utilized them to identify and characterize nine chromosomal loci as centromeres.

In most eukaryotes, the centrochromatin is composed of nucleosomes containing the histone H3 variant CENP-A as a hallmark of centromere identity. The most striking observation of our study was the absence of this centromere determining protein in all Mucoralean species. Absence of CENP-A was accompanied by loss of CENP-C, while most other kinetochore protein orthologs were present in these species. Loss of a previously-thought indispensable protein like CENP-A raised questions on the evolutionary bases of centromere and kinetochore structure. Absence of CENP-A had been previously observed in certain insect orders including butterflies and moths, bugs and lice, earwigs, and dragonflies which in all known cases led to a transition to holocentricity, suggesting that the presence of a centromere-specific protein is dispensable for segregation of a chromosome with localized kinetochore^10^.

Our study is the first to demonstrate loss of core kinetochore proteins in the fungal kingdom, reinforcing that the absence of CENP-A can be tolerated in eukaryotic organisms. Surprisingly, punctate localization patterns of other conserved kinetochore proteins including Mis12, Dsn1, and CENP-T throughout the cell cycle indicate that these organisms still retain amonocentric arrangement of a canonical kinetochore structure, as opposed to the holocentric insects that lack CENP-A. Kinetochore proteins in *M. circinelloides* were observed to be localized at the periphery of each nucleus in the asexual spores as well as mycelia. Three types of nuclear divisions have been studied in multinucleated fungi – synchronous, asynchronous, and parasynchronous^31^. In all ascomycetes except *S. pombe* and *Zymoseptoria tritici*, kinetochore proteins are centromere-localized and clustered throughout the cell cycle^32,33^. In the basidiomycete *C. neoformans*, kinetochore proteins transition between clustering and declustering states in different stages of the cell cycle^34^.

We have observed asynchronous nuclear divisions during germination of asexual spores of *M. circinelloides*, a multinucleated and coenocytic fungus. The kinetochore was found to be consistently present as a cluster in all stages of mitosis, implying that the basic kinetochore structure in this organism is constitutively assembled throughout the cell cycle. Overall, these findings hint that the role of CENP-A is being carried out by either a distinct protein that has a newly evolved function or by one of the known kinetochore proteins like CENP-T, which also has a histone-fold DNA binding domain. We also tried to find whether one of the histone H3 variants plays a role of centromeric histone, but none of them displayed kinetochore-like localization, indicating that they are probably not involved in replacing CENP-A function in chromosome segregation mediated by the kinetochore machinery.

A ChIP-based analysis using Dsn1 and Mis12 revealed the first glimpse of the centromere structure of this Mucoromycotina species. The centromeres in this organism possess features of both point and regional centromeres, resulting in a unique mosaic; thus, we named them mosaic centromeres (Fig. 5). One of their distinctive features is a small kinetochore protein binding domain (∼750-1150 bp) defined by the presence of an AT-rich sequence stretch (∼70-80% AT, ∼100 bp), which is followed by a conserved centromere-specific 41-bp sequence motif. This motif is only found in *M. circinelloides* centromeres and marks the start of the major kinetochore-binding region, suggesting that it could be a binding site for the unknown centromere-specific protein. Overall, this pattern resembles the distribution of conserved elements in *S. cerevisiae* point centromeres, in which two conserved <u>C</u>entromere <u>D</u>etermining <u>E</u>lements (CDEI and CDEIII) are separated by an AT-rich non-conserved DNA sequence stretch (CDEII)^7^.

Other features of *M. circinelloides* mosaic centromeres are reminiscent of fungal regional centromeres like the ones described in *Candida albicans*^35^. The pericentric regions of *M. circinelloides* are large (∼15-75 kb), gene-free, and transcriptionally silenced sequences that are interspersed by Grem-LINE1s, which are repeats of a LINE1-like non-LTR retrotransposable element. These types of retroelements have been identified in the centromeres and neocentromeres of highly diverged eukaryotes, especially LINE1-like elements in mammals^36–41^ and also other non-LTR retroelements as active components of fruit fly centromeres^42^, and LTR retrotransposons in plants^43^ and fungi^22^. Based on these findings, a model has been proposed that CENP-A is recruited by genomic sequences rich in retrotransposable elements and thus, gives rise to the centromeres^44^. *M. circinelloides* is the first organism to display an enrichment in retrotransposable elements in their centromeric regions in the absence of CENP-A, suggesting that these non-LTR elements determine the formation of centromeres. Thus, we hypothesize that identifying Grem clusters in other species of Mucoromycotina could pinpoint the location of their putative pericentromeric regions.

RNAi machinery is essential to control the mobility of transposable elements, but also to maintain centromere structure and stability^22^. Fungi that have lost essential components of the RNAi machinery have shorter pericentric regions and fewer transposable elements than closely related RNAi-proficient species. *M. circinelloides* has a functional and complex RNAi system with Dicer-dependent and -independent pathways^23^. Our results demonstrate that the expression of the centromeric Grem-LINE1s is suppressed by the Dicer-dependent RNAi pathway, which may have resulted in accumulation of these elements in the centromeres^22^.

The composition of centromere-specific nucleosomes has been debated and several conflicting models have been proposed, which include a hemisome in yeast and *Drosophila* with one CENP-A/H4 and one H2A/H2B dimer^45,46^, a conventional octamer with two CENP-A/H4/H2A/H2B^47^, hexasomes with two CENP-A/H4/Scm3, and mixed octasomes which have one molecule of H3 with CENP-A^48^. In addition, H3 nucleosomes have been shown to be interspersed with CENP-A nucleosomes in certain plants and animals^49–51^. We observed the presence of H3 at the relatively short kinetochore protein-binding region at a level that is similar to the pericentric region or gene bodies. Because histone H3 was found to be enriched at the core centromere, we propose that either a homotypic histone H3 nucleosome or a heterotypic histone H3 nucleosome exists where CENP-A is replaced with some other protein. Further studies will provide the exact composition of centromere-specific nucleosomes in this basal fungus.

Overall this study has uncovered for the first time the centromeres of an early-diverging fungus, which are characterized by a mosaic of features from both point and regional centromeres found in the fungal kingdom. Several studies on rapidly-evolving centromeres suggest that point centromeres may be a recently evolved state as compared to regional centromeres^9^. Therefore, we hypothesize that the last mucoralean common ancestor had regional centromeres with pericentric regions that harbored retrotransposable elements contained by an active RNAi machinery. The loss of CENP-A triggered an evolutionary rearrangement from typical regional centromeres to the smaller kinetochore-binding regions that defines the mosaic centromeres of *M. circinelloides*.

## Methods

### Fungal strains and culture conditions

All the strains used and generated in this work derive from *M. circinelloides* f.*lusitanicus* CBS277.49. The double auxotroph MU402^14^ (Ura^−^, Leu^−^) or the MU636 (Ura^+^, Leu^−^) strain was used for transformation with cassettes containing the selectable marker for uracil *pyrG* or for leucine *leuA*, respectively. Single tagged strains in kinetochore proteins MU840, MU842, MU844, and MU846 were used as parental strains for transformation with the Hht4-eGFP cassette to obtain double-tagged strains. All the generated strains in this work are listed in Supplementary Table 3.

After transformation, to select uracil prototrophy *M. circinelloides* spores were plated in minimal medium with Casamino Acids (MMC)^14^, and for leucine prototrophy spores were plated in medium yeast nitrogen base (YNB)^14^, both adjusted at pH 3.2 for transformation and colony isolation. *M. circinelloides* spores were cultured in rich medium yeast-peptone-glucose (YPG)^14^ for optimal growth and sporulation at pH 4.5. The microscopy and ChIP assays were performed with pre-germinated spores growing in YPG pH 4.5 for 3 hours at 250 rpm and 26 ° C.

### Microscopy imaging and acquisition

Freshly isolated *M. circinelloides* spores were washed twice with 1X phosphate buffered saline (PBS). The spores were resuspended in PBS and 10 μl suspension was placed on a slide, covered with a coverslip and sealed with clear nail polish. The images were acquired at room temperature using laser scanning inverted confocal microscope LSM 880-Airyscan (Zeiss, Plan Apochromat 63x, NA oil 1.4) equipped with highly sensitive photo-detectors. The filters used were GFP/FITC 488, mCherry 561 for excitation and GFP/FITC 500/550 band pass, mCherry 565/650 long pass for emission. Z-stack images were taken at every 0.5 μm and processed using Zeiss Zen software. All the images were displayed after the maximum intensity projection of images using Zeiss Zen software.

### Live cell imaging

Freshly isolated *M. circinelloides* spores were incubated in YPG growth medium at 26°C for 2 h. Then, the spores to be imaged were pelleted at 3000 xg and washed twice with 1X phosphate buffered saline (PBS). The spores were resuspended in PBS and 10μl suspension was placed on a slide containing thin 2% agarose patch containing 2% dextrose and covered with a coverslip. Live cell imaging was performed at 30°C in a temperature-controlled chamber of an inverted confocal microscope (Zeiss, LSM-880) using a Plan Apochromat 100X NA oil 1.4 objective and GaAsp photodetectors. Images were collected at 60-s intervals with 0.5 µm Z-steps using GFP/FITC 488, mCherry 561 for excitation. GFP/FITC 500/550 band pass or GFP 495/500 and mCherry 565/650 or mCherry 580-750 band pass for emission. All the images were displayed after the maximum intensity projection of images at each time using Zeiss Zen software.

### Ortholog search

Seventy five fungal proteomes were retrieved from the Joint Genome Institute (JGI) Mycocosm genome portal^52^ (Supplementary Table 1). BLAST+^53^ v2.7.1 individual protein-protein BLASTp, iterative BLASTp (PSI-BLAST) and iterative HMMER v3.2.1 (http://www.hmmer.org) jackhammer searches (E-value ≤ 1×10^−5^) were conducted against these proteomes using *M. musculus, U. maydis, S. cerevisiae*, and *S. pombe* protein sequences (Supplementary Table 1) as queries. In addition, an hmmsearch was launched using the Pfam-A^54^ HMM models for each kinetochore protein [Pfam gathering threshold (GA) as cut-off value]. All the putative orthologs found were used to perform a reciprocal BLASTp against *M. musculus, U. maydis, S. cerevisiae*, and *S. pombe* protein sequences. Also, Pfam-A protein domains were searched in all the possible orthologs using HMMER hmmscan. Protein sequences that either lacked appropriate Pfam domains or failed to produce a hit in a reciprocal blastp search were discarded, resulting in a matrix of putative orthologs for each given species (Supplementary Table 1). The cladogram showing the relationship among species was drawn using the tree data from JGI Mycocosm^52^ and the interactive <u>T</u>ree <u>o</u>f <u>L</u>ife^55^ (iTOL v4) tool. To determine CENP-A presence or absence in each given species, all 289 protein sequences of histone H3 putative orthologs (Supplementary Table 1) were aligned using MUSCLE^56^ v3.8.1551, and the phylogenetic relationship of their HFD was inferred by the neighbor-joining method employing the Jones, Taylor, and Thornton (JTT) substitution model^57^, with a bootstrap procedure of 1,000 iterations (MEGA^58^ v10.0.5).

### Construction of the strains expressing the histone and kinetochore proteins with the fluorescent tag

Alleles containing fluorescent fusion protein coding-sequences were designed to either integrate them in their corresponding endogenous locus, or in the *carRP* locus. The Hht4-eGFP, Hhf1-eGFP, Hhf4-eGFP and CENP-T-eGFP alleles contained the ORF of each gene fused in-frame at their C-termini with the eGFP sequence, followed by the *leuA* gene as a selectable marker. The alleles were obtained by overlapping PCR using 5’-modified primers harboring restriction sites for easy cloning. Briefly, each allele was obtained by fusing the 1-kb fragment upstream the stop codon of each gene, the 0.7-kb eGFP fragment (ending in a stop codon), the 3.4-kb *leuA* fragment, and the 1-kb sequence downstream the stop codon of each tagged gene. All the linear alleles were digested with appropriate restriction enzymes, ligated into the plasmid Bluescript SK (+), and cloned into *Escherichia coli*.

The Mis12-mCherry and Dsn1-mCherry constructions were designed to allow the integration and ectopic overexpression of the fused proteins in the *carRP* locus. First, a universal mCherry-expression vector was constructed, containing the strong promoter *Pzrt1*, the ORF of the mCherry coding-gene including the stop codon, and the *pyrG* gene as a selectable marker (named pMAT1915). To do that, a fragment containing the 1-kb downstream and upstream sequences from the *carRP* gene flanking the *Pzrt1* promoter was amplified with an inverse PCR using the plasmid pMAT1477^14^ as a template. Then, the *mCherry* and *pyrG* fragments were added by In-Fusion cloning (Takara) using primers with appropriate overlapping 5’-tails (Supplementary Table 2). Once the universal pMAT1915 vector was constructed, it was used for the C-terminal tagging of *mis12* or *dsn1*, fusing their ORFs without the stop codon upstream and in-frame with the *mCherry* ORF by In-Fusion cloning. The resulting plasmids were pMAT1917 and pMAT1919, respectively for *mis12* and *dsn1*. For the N-terminal tagging, the plasmid pMAT1915 was inverse-amplified using primers that exclude the stop codon of the *mCherry* fragment, and the *mis12* and *dsn1* ORFs, including their stop codons, were cloned in-frame. The resulting plasmids were pMAT1916 and pMAT1918 for mCherry-Mis12 and mCherry-Dsn1 expression, respectively.

Stellar™ competent cells (Takara) were used for transformation by heat-shock following the supplier procedures. The correct construction of all the generated plasmids was confirmed by restriction analysis and Sanger sequencing.

For transformation, the alleles were released from the vector backbone by restriction digestion and used to transform *M. circinelloides* protoplasts by electroporation^14^. Spores from individual colonies were collected and plated in selective media to ensure homokaryosis. After at least five vegetative cycles, the integration in the target locus was confirmed by PCR using primers that amplified the whole locus (Supplementary Table 2), producing PCR-fragments that allowed discrimination between the mutant and the wild-type locus (Supplementary Fig. 3, 4). Genomic DNA purification was performed following the procedure previously published^59^.

### Chromatin immunoprecipitation and sequencing

The ChIP protocol for *M. circinelloides* was adapted from previous studies in other fungal models^22^. Briefly, spores from the strains MU842 (expressing Mi12-mCherry) and MU846 (expressing Dsn1-mCherry) were used for independent IP experiments. 10^7^ spores/mL of these strains were germinated in 100 ml of liquid YPG pH 4.5 medium, shaking at 250 rpm at 26°C for 3 hours. Then, cells were fixed by adding formaldehyde at a final concentration of 1% for 20 minutes. The fixation was quenched by adding glycine to a final concentration of 135 mM. Cells were pelleted and ground with liquid nitrogen, and 150 mg of ground pellet was resuspended in ChIP lysis Buffer (50 mM HEPES pH 7.4, 150 mM NaCl, 1% Triton X-100, 0.1% DOC, and 1 mM EDTA), with a protease inhibitor cocktail. The sample was sonicated using a Bioruptor (Diogenode) with pulses of 30s ON/OFF for 50 cycles to obtain a chromatin fragmentation between 100 and 300 bp.

The lysates were clarified by centrifugation at 12,000 xg for 10 min at 4°C. At this step, 100 μL were frozen at −20°C for Input DNA control. The samples were then divided into 450 μL for the antibody immunoprecipitation and 450 μL for the mock control. For *IP* 25 μL of <u>R</u>ed <u>F</u>luorescent <u>P</u>rotein (RFP)-Trap <u>M</u>agnetic <u>A</u>garose (MA) beads (Chromotek) were added and 25 μL of MA beads (Chromotek) to the binding control. Samples were incubated at 4°C overnight. After incubation, the beads were washed twice with 1 mL of the low salt wash buffer (20 mM Tris pH 8.0, 150 mM NaCl, 0.1% SDS, 1 % Triton X-100, and 2 mM EDTA), twice with 1 mL of the high salt wash buffer (20 mM Tris pH 8.0, 500 mM NaCl, 0.1% SDS, 1% Triton X-100 and 2 mM EDTA), once with the LiCl wash buffer (10 mM Tris pH 8.0, 1 mM EDTA, 0.25 M LiCl, 1% NP40, and 1% DOC), and finally once with TE (10mM Tris pH 8.0, and 1 mM EDTA). To elute the DNA, 250 μL of TES (50 mM Tris pH 8.0, 10 mM EDTA, 1% SDS) was added to each sample and incubated for 10 min at 65°C, twice. The immunoprecipitated samples and the Input controls were de-crosslinked by incubating at 65°C overnight. The samples were treated with 199 μg of RNase A (Sigma Aldrich) and 190 μg of Proteinase K (Sigma Aldrich) for 2 hours at 50°C. For DNA purification, 1 volume of phenol:chloroform:isoamyl alcohol (25:24:1) was added, the aqueous phase was recovered, and then 1 volume of chloroform:isoamyl alcohol (24:1) was added. After centrifugation, the aqueous phase was recovered, and DNA was precipitated with 20 μg of glycogen (Thermo Fisher), 1/10 volume of Na-Acetate 3 M (pH 5.2), and 1 volume of ethanol 100%. Pulled DNA from duplicated samples were precipitated together for sequencing. Samples incubated at −20°C overnight were centrifuged for 10 min at 12,000 xg and the pellets were washed with ethanol 70%. Each sample was resuspended in 30 μL of MilliQ water. The quality of the DNA was analyzed by QuBit (Thermo Fisher) and sent to Novogene, which prepared the libraries using TruSeq ChIP Library Prep Kit and sequenced them with Illumina Hiseq 2500 High-Output v4 single-end reads of 50 bp.

Histone H3-ChIP assays were performed in the same way with the following changes. The protoplasts were obtained as described before^60^ and then sonicated as above. Native ChIP was performed^22^ using anti-H3 antibody (Abcam ab1791). The DNA pellet was dissolved in 25 μl of MilliQ water. All samples (*Input, IP* with or without antibodies) were used for PCR. The *Input* and *IP* DNA was subsequently used for qPCR using the primers listed in Supplementary Table 2b. Histone H3 enrichment was determined by the percentage input method using the formula:100 *x* 2^*adjusted Ct Input*−*adjusted Ct IP*^ (ref). The adjusted Ct value is the dilution factor (log2 of dilution factor) subtracted from the Ct value of the *Input* or *IP* DNA samples. Three technical replicates were taken for qPCR analysis and standard error of mean was calculated.

### Sequencing data analysis

Raw reads from <u>Ch</u>romatin Immunoprecipitation and <u>RNA</u> <u>seq</u>uencing (ChIP- and RNA-seq) were quality-checked with FASTQC v0.11.8 and adapters removed with Trim Galore! v0.6.2 (http://www.bioinformatics.babraham.ac.uk/projects/). Reads were aligned to the *Mucor circinelloides* f. *lusitanicus* MU402 reference genome Muccir1_3 (referred to as the *M. circinelloides* genome throughout the text, available at JGI http://genome.jgi.doe.gov/Muccir1_3/Muccir1_3.home.html), using the Burrows-Wheeler Aligner^61^ (BWA v0.7.8) algorithm for ChIP- and sRNA-seq reads and STAR^62^ v2.7.1a for mRNA-seq data. The number of overlapping aligned reads per 25-bp bin was used as a measure of coverage, obtaining normalized bigWig coverage files with deepTools^63^ v3.2.1 bamCoverage function. For RNA-seq data, coverage was normalized to bins per million mapped reads (BPM). For ChIP-seq data, reads were normalized per genomic content (1x normalization). ChIP enrichment peaks were identified by <u>M</u>odel-based <u>A</u>nalysis of <u>C</u>hIP-<u>s</u>eq^64^ (MACS2 v2.1.1) callpeak function and sorted by fold-enrichment and FDR values (Supplementary Table 4). Enriched peaks (fold-enrichment ≥ 1.6, FDR ≤ 5×10^−5^) were manually curated by analyzing the *IP* coverage with the Integrative <u>G</u>enomics <u>V</u>iewer^65^ (IGV v2.4.1) genome browser, ensuring that peaks were present in both Mis12 and Dsn1 *IP* samples, and absent in either the input and binding controls. The resulting centromeric sequences were analyzed with <u>M</u>ultiple <u>E</u>m for <u>M</u>otif <u>E</u>licitation^66^ (MEME v5.0.5) to discover conserved DNA motifs on both strands. GC content every 25-bp bin was calculated with bedTools^67^ v2.28. Normalized coverage, GC content and genetic elements annotation files were rendered with either Gviz^68^ v1.29.0 for whole-chromosome visualization or deepTools pyGenomeTracks module for close-in genomic views.

### Transposable element analysis

The *M. circinelloides* genome assembly of strain MU402 was searched for repeats using RepeatScout^69^ v1.0.5 and RepeatMasker (http://www.repeatmasker.org) v4.0.9. The repeats located in the pericentromeric regions were aligned using MAFFT^70^ v7.407 and clustered with CD-HIT^71^ v4.8.1, generating groups of sequences that shared ≥ 95% identity and ≥ 70% coverage and calculating their *p*-distance (nucleotide changes per site). A BLASTn search against *M. circinelloides* genome assembly was conducted using a representative repeated sequence from each cluster to locate similar sequences across the entire genome, obtaining hits with ≥ 80% identity and ≥ 20% coverage (Supplementary Table 5). The pericentromeric repeats were examined with getorf from Emboss^72^ v6.6.0, obtaining all ORFs ≥ 600 nt. Protein domains in these ORFs translated sequences were predicted by hmmsearch using the Pfam-A v32.0 database. The reverse transcriptase (RVT) domains found in the repeated sequences were aligned with MUSCLE to create a consensus sequence, and this consensus sequence was aligned with 281 RVT domains of well-known non-LTR retrotransposons from Repbase^73^ 24.04. A neighbor-joining phylogenetic tree was inferred from this alignment using the JTT substitution model and a bootstrap procedure of 1000 iterations (MEGA v10.0.5). To assess the presence of the retrotransposable elements in other species of Mucoromycotina, a tBLASTn search was launched against the 55 available genomes at JGI (Supplementary Table 6) using the translated sequence of both ORFs combined, recovering hits with ≥ 50% coverage and E-value ≤ 10^−5^.

## Data availability

ChIP-seq raw data and processed files are deposited at the <u>G</u>ene <u>E</u>xpression <u>O</u>mnibus (GEO) repository under GSE132687 accession number. Open access will be granted upon publication and they are provisionally available upon token request. Both mRNA and sRNA-seq data were already available at the Sequence Read Archive (SRA) and can be accessed with the following run accession numbers: SRR1611144 (R7B wild-type strain mRNA), SRR1611151 (*ago1* deletion mutant strain mRNA), SRR1611171 (double *dcl1 dcl2* deletion mutant strain mRNA), SRR039123 (R7B wild-type strain sRNA), SRR836082 (*ago1* deletion mutant strain sRNA), SRR039128 (double *dcl1 dcl2* deletion mutant strain sRNA).

## Acknowledgments

This work was supported by the Ministerio de Economía y Competitividad, Spain (grant number BFU2015-65501-P, cofinanced by FEDER, and RYC-2014-15844) and Ministerio de Ciencia, Innovación y Universidades, Spain (grant number PGC2018-097452-B-I00, cofinanced by FEDER). CPA and MINM were supported by predoctoral fellowships from the Ministerio de Educación, Cultura y Deporte, Spain (grants number FPU14/01983 and FPU14/01832 respectively). We acknowledge financial support from Tata Innovation Fellowship, Department of Biotechnology, Government of India (BT/HRD/35/01/03/2017), DBT-BUILDER programme support at JNCASR (BT/INF22/SP27679/2018) and intramural support from JNCASR to KS. SP is supported by DST-WOS-A postdoctoral fellowship (SR/WOS-A/LS-207/2018). Supported in part by NIH/NIAID R37 grant AI39115-21 and R01 grant AI50113-15 to J.H., and by the CIFAR program Fungal Kingdom: Threats & Opportunities, for which J.H. serves as Co-Director and Fellow. We thank B. Suma at the confocal imaging facility of the Molecular Biology and Genetics Unit, JNCASR for assistance in imaging. Work conducted by the US Department of Energy Joint Genome Institute, a DOE Office of Science User Facility, was supported by the Office of Science of the US Department of Energy under Contract No. DE-AC02-05CH11231. We thank Alexander Idnurm (University of Melbourne, Australia), Luis M. Corrochano (University of Seville, Spain), Timothy Y. James (University of Michigan, USA), Mathew E. Smith (University of Florida, USA), Joseph W. Spatafora (Oregon State University, USA), Jason E. Stajich (University of California-Riverside, USA), and Gregory Bonito (Michigan State University, USA) for sharing fungal genome sequences to analyze the distribution of kinetochore proteins. We also thank Thomas D. Petes (Duke University School of Medicine, USA) and Beth A. Sullivan (Duke University School of Medicine, USA) for critical reading of the manuscript and the members of KS lab, VG lab, and JH lab for valuable discussions during bi-weekly Skype meetings.

## Author contributions

KS, JH, VG, and FEN conceived and supervised the study. MINM and CPA analyzed the distribution of kinetochore proteins in fungi, generated and validated the plasmids and strains, performed the ChIP-seq experiments, identified and defined the retrotransposable elements, and analyzed the mRNA and sRNA-seq data. MINM, CPA and PG analyzed the ChIP-seq data. MINM, CPA, PG and SP characterized the centromeric loci. SP, MINM and CPA did the fluorescence microscopy imaging. SP performed the ChIP-qPCR experiments. IVG, JP and SJM provided the genome assembly and gene annotation. MINM, CPA and SP designed the figures and wrote the manuscript with the support from KS, JH, and VG. VG, JH, FEN and KS provided funding. All authors revised the manuscript and approved the final version.

---

**Supplementary Fig. 1. Mucorales and Umbelopsidales lack CENP-A. (a)** Schematic of canonical histone H3 and CENP-A shared protein features. **(b-d)** Multiple protein sequence alignments of the N-tail **(b)**, Loop 1 **(c)**, and C-terminal **(d)** regions. The scale above the comparisons indicates the amino acid positions with *S. pombe* sequence as the reference. Red arrows mark relevant amino acid positions. Amino acids are colored in increasing shades of blue to show conservation within each group. **(e)** Neighbor-joining phylogenetic tree (JTT model) of the <u>H</u>istone <u>F</u>olding <u>D</u>omain (HFD), showing phylogenetic distance (branch length) and branch support (1000 bootstraps). Branches with < 50% bootstrap support are collapsed. The protein sequences analyzed in **b** to **e** are distributed among three groups: well-studied histone H3 (yellow), rare histone H3 (blue), and the predicted CENP-A proteins in this study (red). Asterisks (*) indicate proteins from the Mucoromycotina.

**Supplementary Fig. 2. *M. circinelloides* histones H3 and H4 are not centromere-specific binding proteins. (a)** Location of all the histone H2A, H2B, H3, and H4-coding genes in the M. *circinelloides* genome. Genes are represented by arrows showing transcription direction; the coding sequence is depicted as larger red blocks, flanked by smaller blue blocks representing the untranslated regions, and connected by black lines as intronic sequences. Light-gray arrows indicate neighboring non-histone genes and their transcription direction. **(b, c)** Protein alignment of several well-characterized H3 **(b)** and H4 **(c)** histones with *M. circinelloides* orthologs. A scale indicates the amino acid positions taking *S. pombe* sequence as the reference. <u>H</u>istone folding <u>d</u>omains (HFD) are outlined in a diagram below each alignment, as well as the N-terminal tail. Amino acids are colored in increasing shades of blue and a consensus protein logo is provided to reflect conservation. **(d)** Confocal microscopy images of *M. circinelloides* strains expressing eGFP-fluorescent fusion histone proteins Hht4, Hhf1, and Hhf3 in 4-hour pregerminated spores. The fluorescent signal is colored as green. A calibrated scale (white bar) is provided for size comparison (5 µm).

**Supplementary Fig. 3. Generation of strains expressing fluorescent fusion histones**. Diagram of the fluorescent fusion alleles used for integration by homologous recombination (shown as discontinuous crosses) into the wild-type allele of each histone loci: *hht4* **(a)**, *hhf1* **(b)**, and *hhf3* **(c)**. Arrows in the diagrams mark the annealing regions for specific primers used to confirm the integration by PCR, amplifying both the wild-type (WT) and mutant (MUT) alleles. Gel images below the diagrams show the size-specific PCR fragments. M lanes were loaded with the GeneRuler DNA Ladder Mix (Thermo Fisher) to estimate the fragment sizes. Green regions indicate mCherry and eGFP genes.

**Supplementary Fig. 4. Generation of mutant strains expressing fluorescent kinetochore proteins**. Diagram (above) and agarose gel images (below) showing the integration of the *cnpT* fusion allele into its wild-type locus **(a)**; and the *mis12* **(b)** and *dsn1* **(c)** fusion alleles (both N-terminal and C-terminal fusions) into the *carRP* locus. Discontinuous crosses indicate homologous recombination and arrows mark the annealing regions for specific primers used to confirm the integration by PCR, amplifying both the wild-type (WT) and mutant (MUT) alleles shown in the gel images. M lanes were loaded with the GeneRuler DNA Ladder Mix (Thermo Fisher) to estimate the fragment sizes. Red and green regions indicate mCherry and eGFP genes, respectively

**Supplementary Fig. 5. *M. circinelloides* has nine centromeres**. *IP* enrichment (coverage x1) of immunoprecipitated DNA (*IP* DNA) from Mis12 and Dsn1 mCherry-tagged strains compared to their corresponding input (*Input* DNA) and binding controls (*beads only* DNA), shown as color-coded tracks across the whole sequence of scaffolds 01-11. Asterisks (*) mark significant peaks in both *IP* DNA samples that are not present in the controls, indicating the putative centromeric regions.

**Supplementary Fig. 6. *M. circinelloides* pericentromeric regions**. A genomic view of the pericentromeric regions of all nine centromeres. Each region shows the kinetochore-binding region enrichment (IP, an average of both *IP* signals minus *Input* and *Beads only* controls), annotated genes (red blocks) and transposable elements (light blue blocks), *CEN*-specific DNA motif position (black vertical line) and its direction (arrow), GC content, and transcriptomic data of mRNA (green) and sRNAs (red) in *M. circinelloides* wild-type strain, and *ago1* and double *dcl1 dcl2* deletion mutants after 48 h of growth in rich media.

**Supplementary Fig. 7. Centromere-specific non-LTR L1-like retrotransposable elements. (a)** Matrix showing pairwise *p*-distance (% changes per site) across all 44 repetitive elements flanking the centromeres of *M. circinelloides*. Elements sharing ≥ 95.0% identity are clustered together, generating 10 groups that comprise full-length elements and incomplete sequences (remnants). **(b)** Neighbor-joining phylogeny (JTT model) of the RVT domain of well-known non-LTR elements. The RVT domain of the centromere-specific element (Grem LINE1) identified in this study is marked inside a blue box. Phylogenetic distance (branch length) and branch support (1000 bootstraps, size-coded circles) are shown. Leaves within the same clade of non-LTR retrotransposons are collapsed, forming a scalene triangle (top and bottom corners show the shortest and longest phylogenetic distance, respectively); except for the L1 clade which contains the Grem LINE-1.

