## Supplementary Figures for "Early diverging fungus *Mucor circinelloides* lacks centromeric histone CENP-A and displays a mosaic of point and regional centromeres"

### Supplementary Figure 1

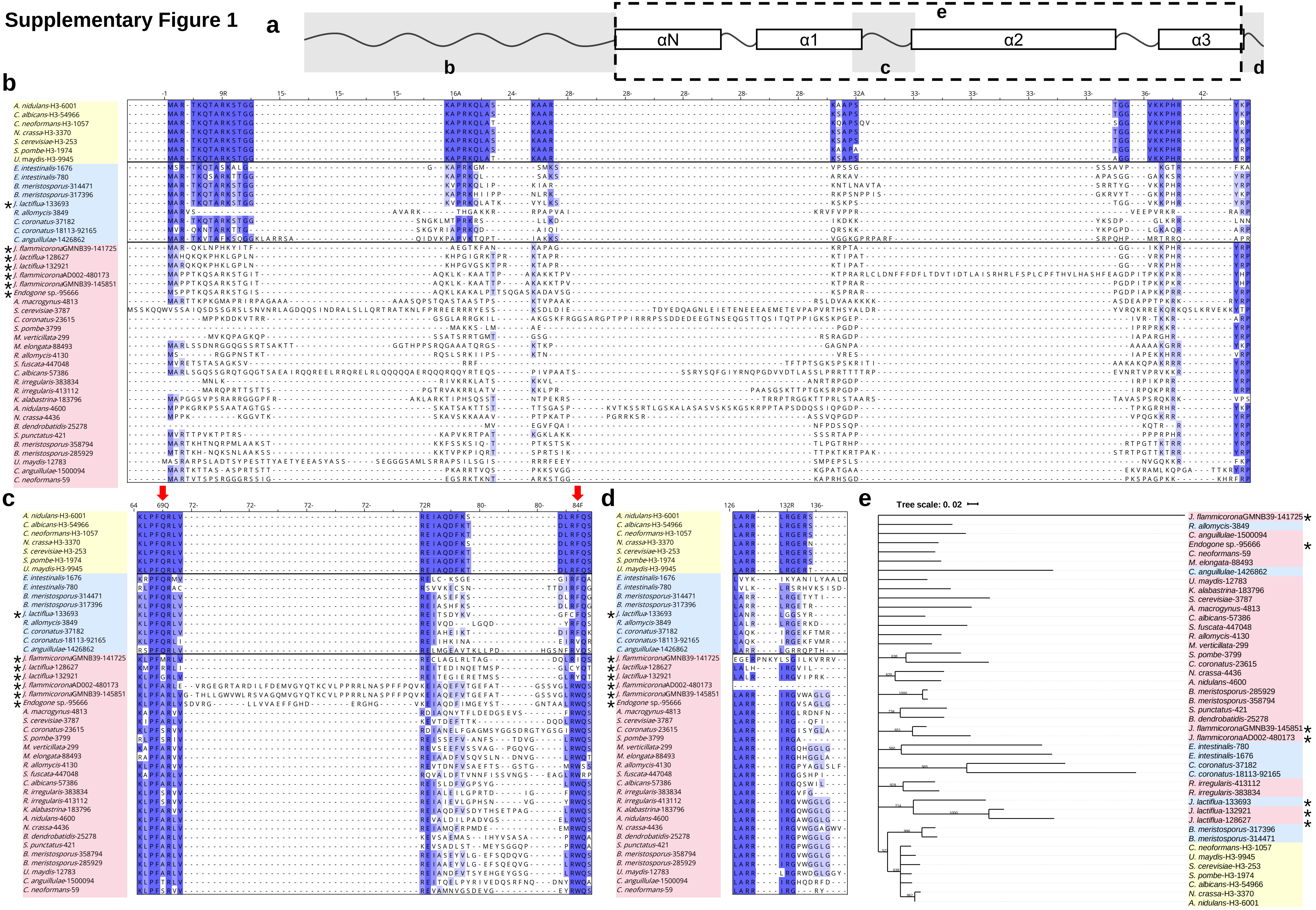

#### Supplementary Figure 2

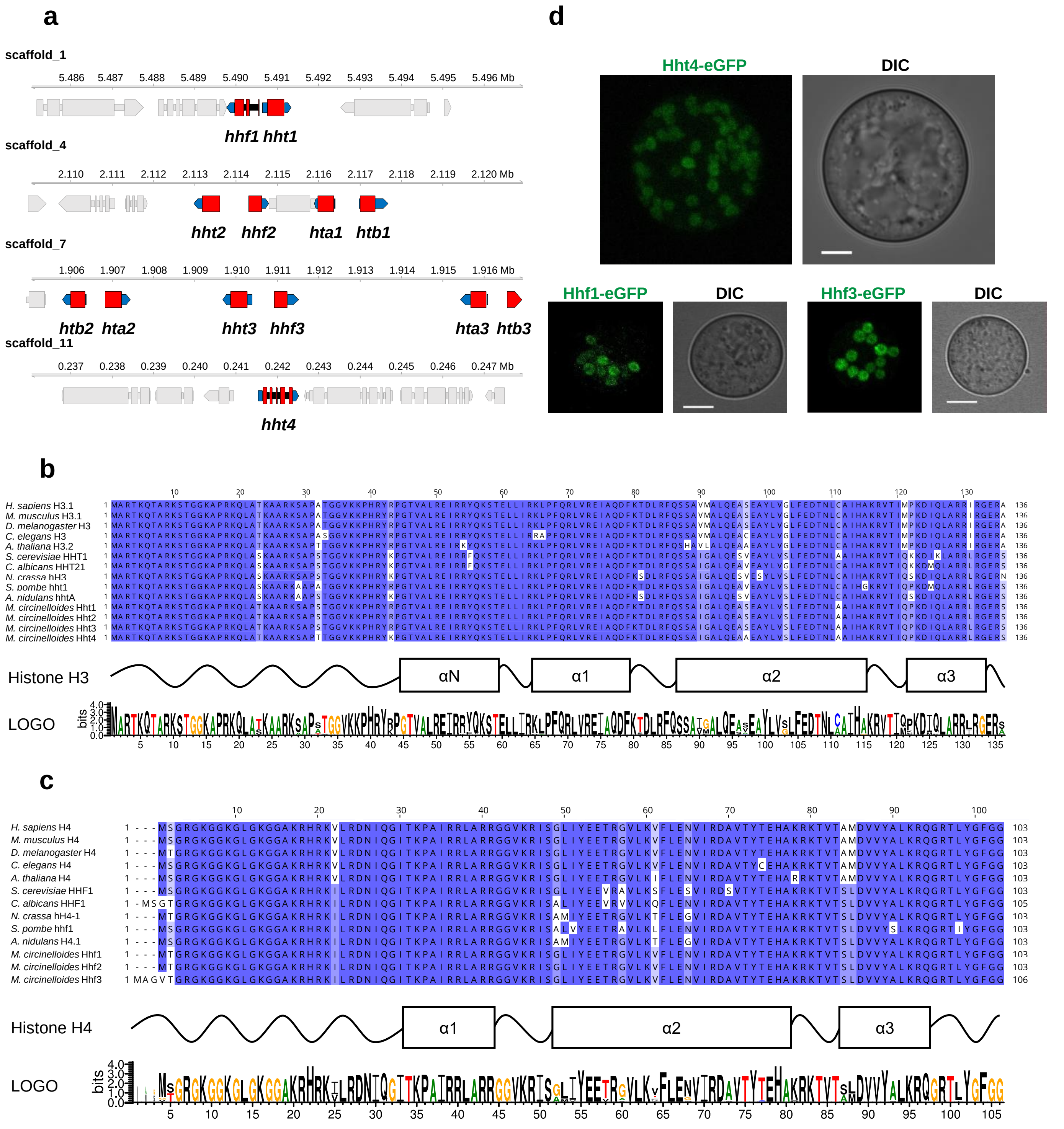

### Supplementary Figure 3

a

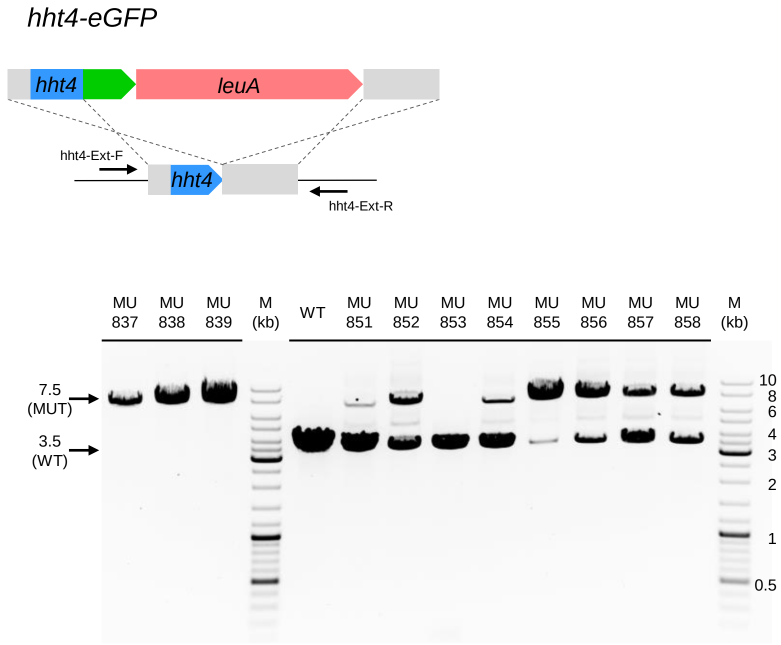


b

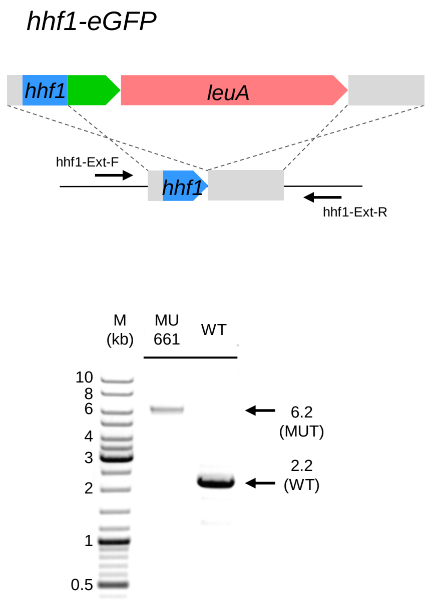


c

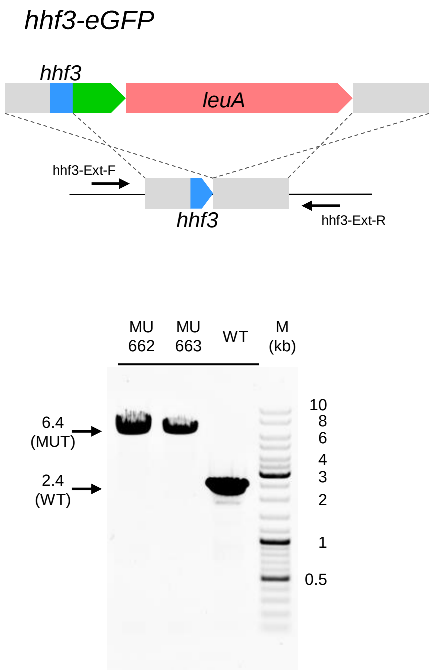

### Supplementary Figure 4

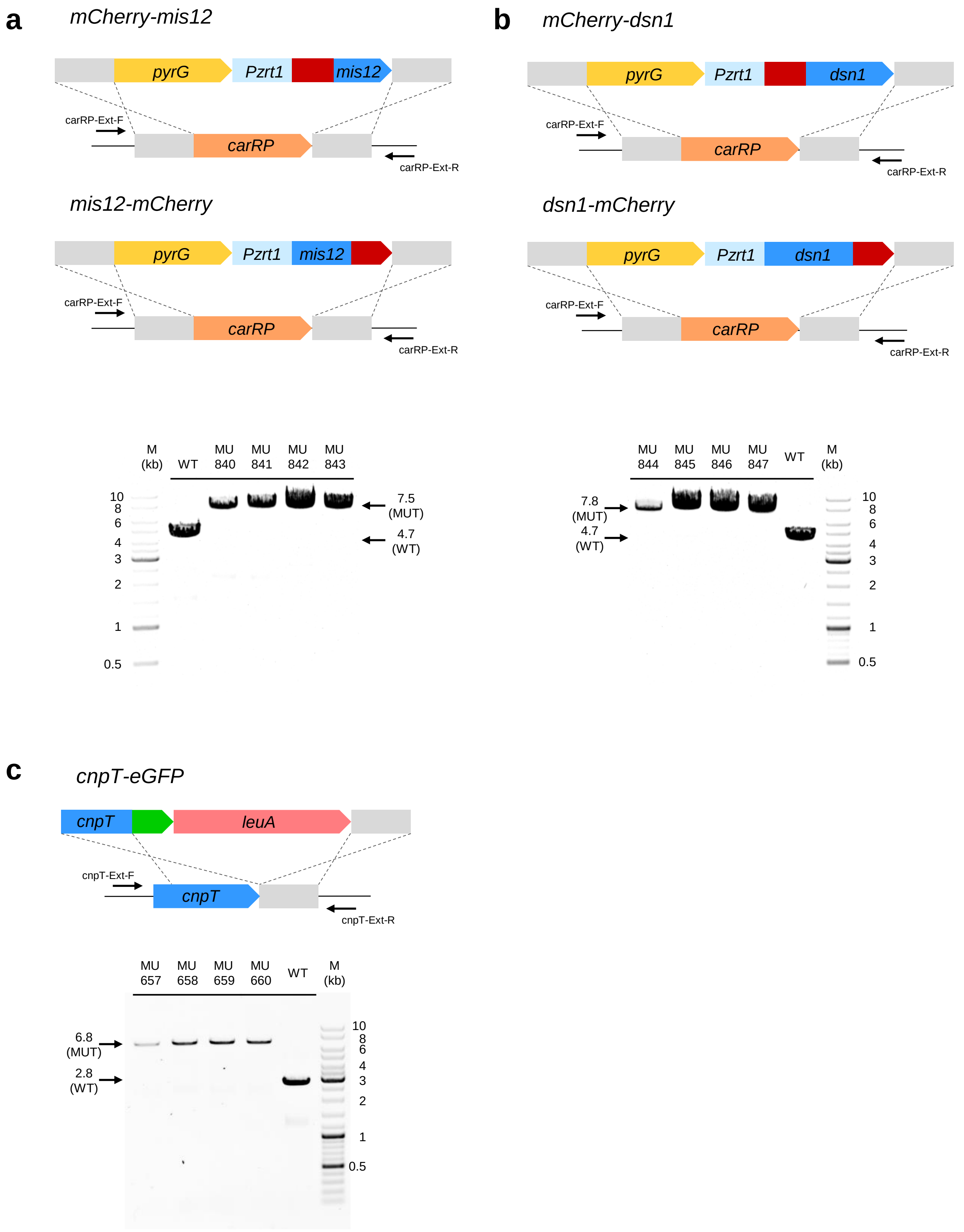

### Supplementary Figure 5

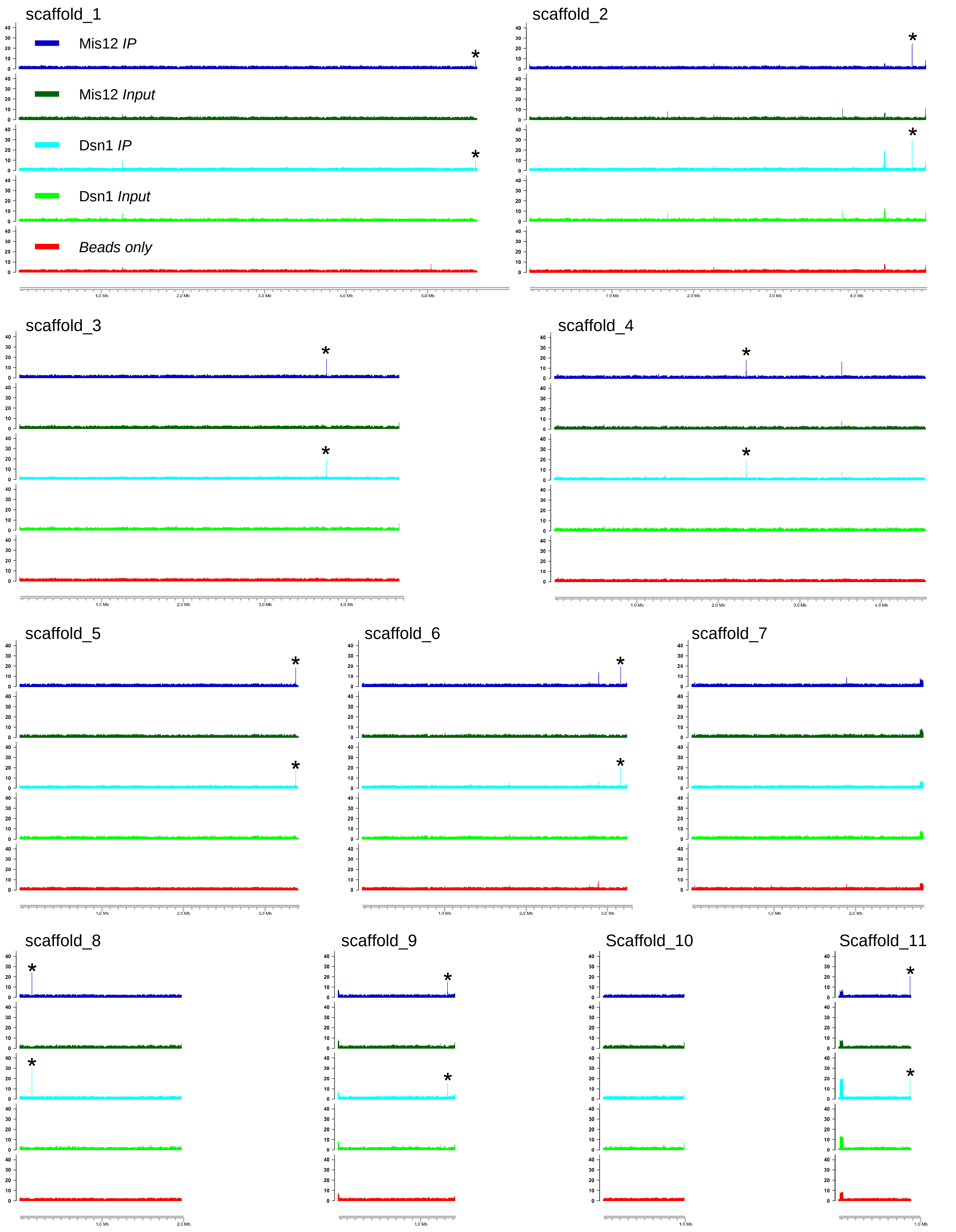

### Supplementary Figure 6

#### *CEN1*

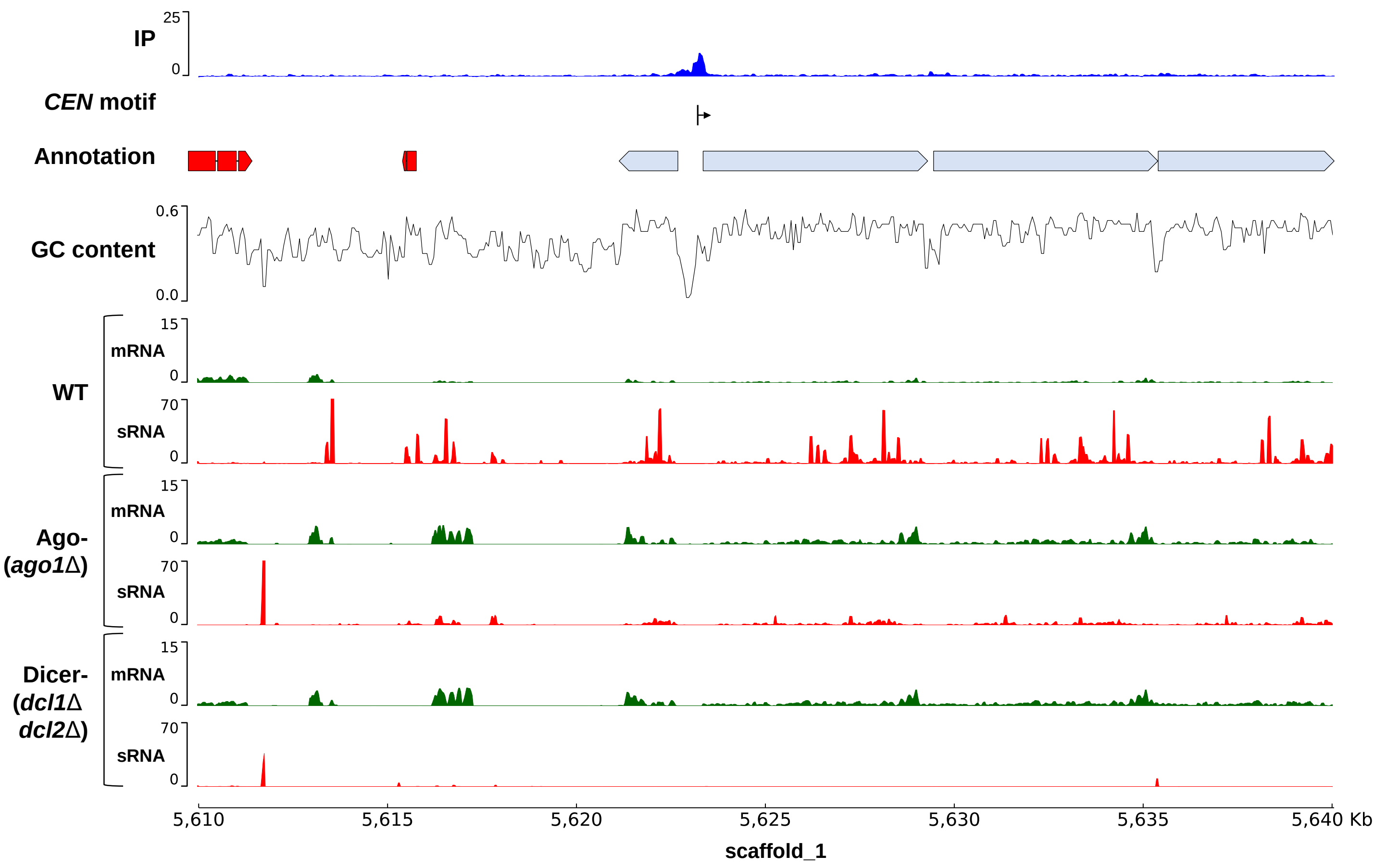


#### *CEN2*

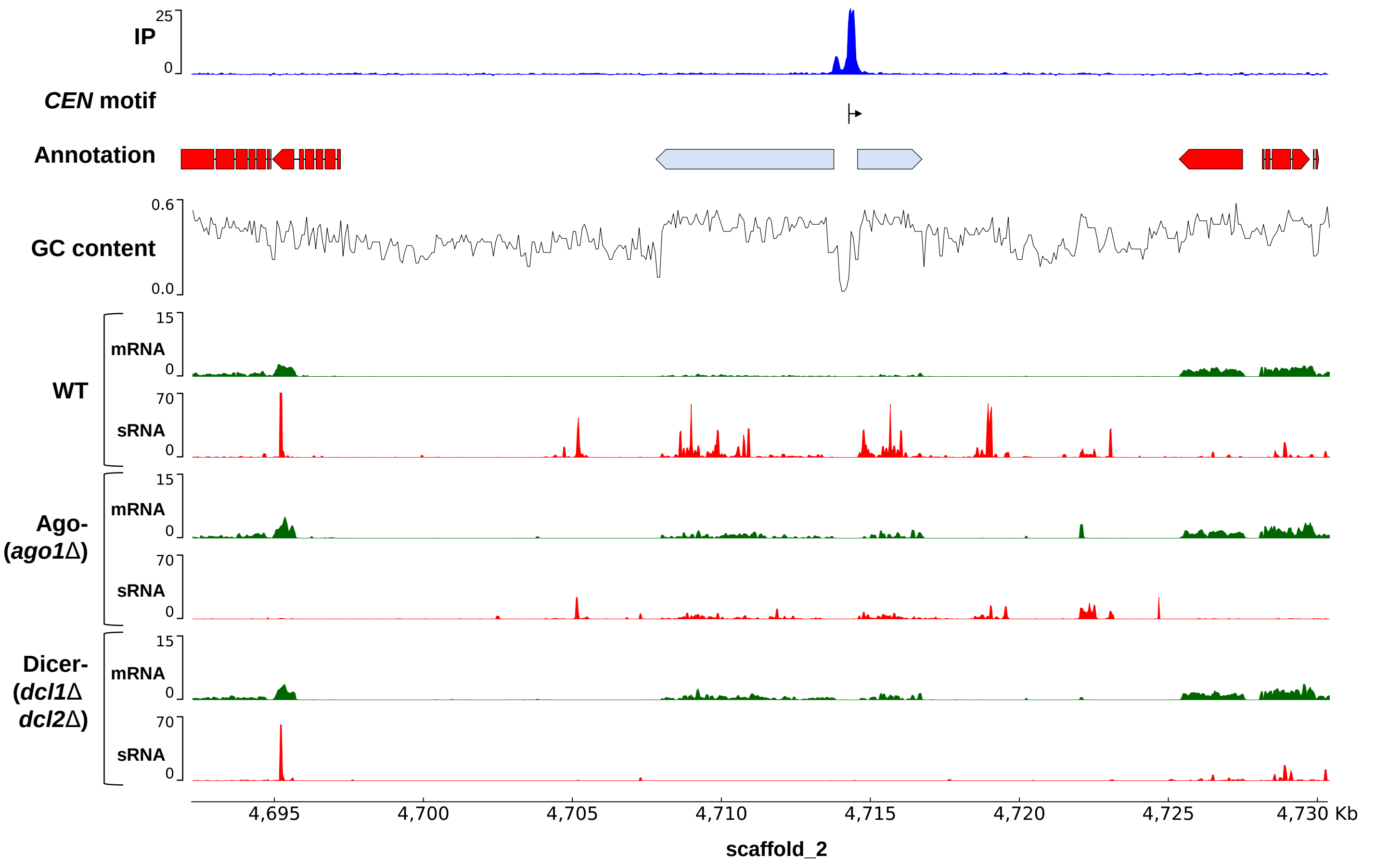

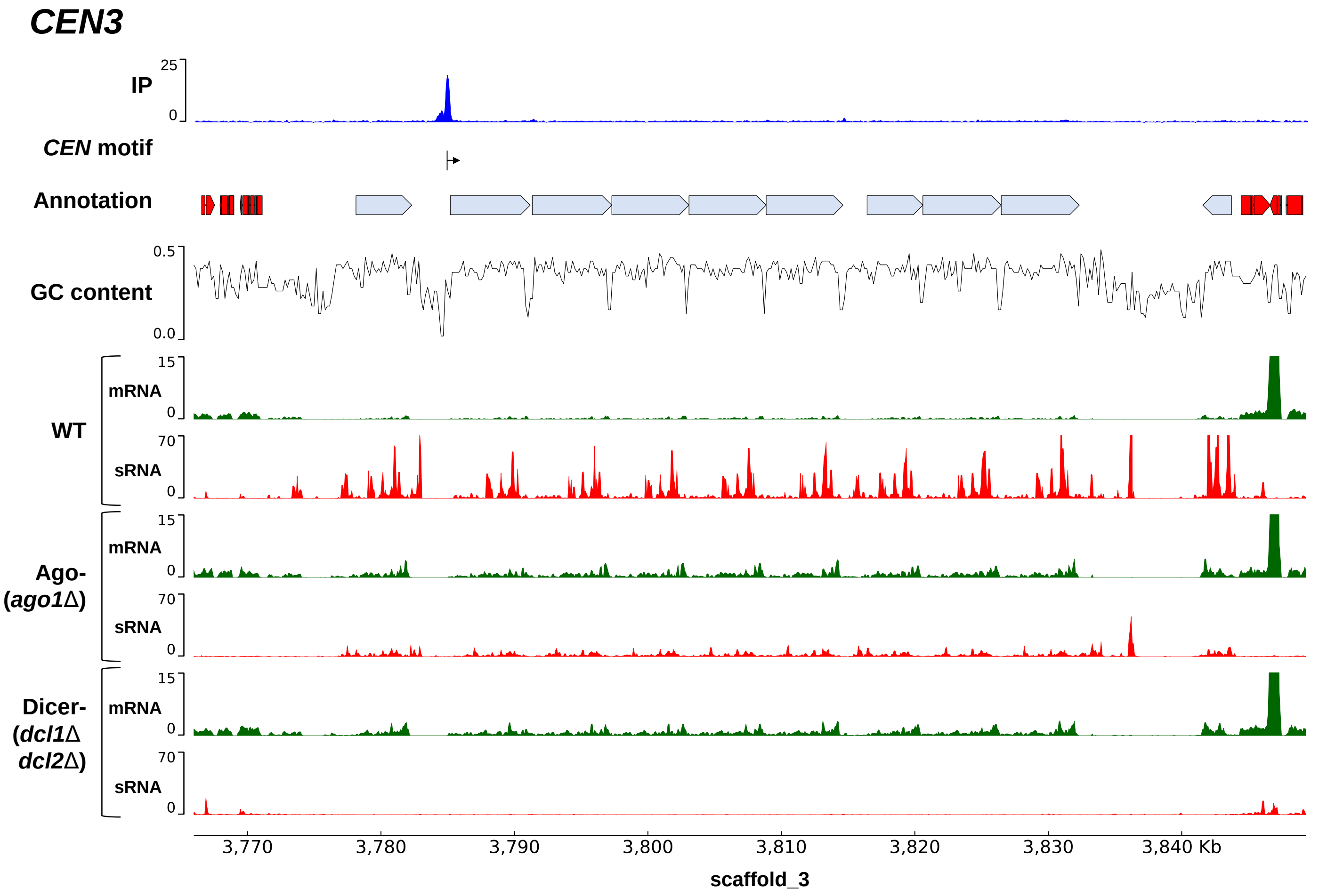
CEN3
IP
25
0
CEN motif
Annotation
GC content
0.5
0.0
WT
mRNA
15
0
sRNA
70
0
Ago-
(ago1Δ)
mRNA
15
0
sRNA
70
0
Dicer-
(dcl1Δ
dcl2Δ)
mRNA
15
0
sRNA
70
0
3,770
3,780
3,790
3,800
3,810
3,820
3,830
3,840 Kb
scaffold_3

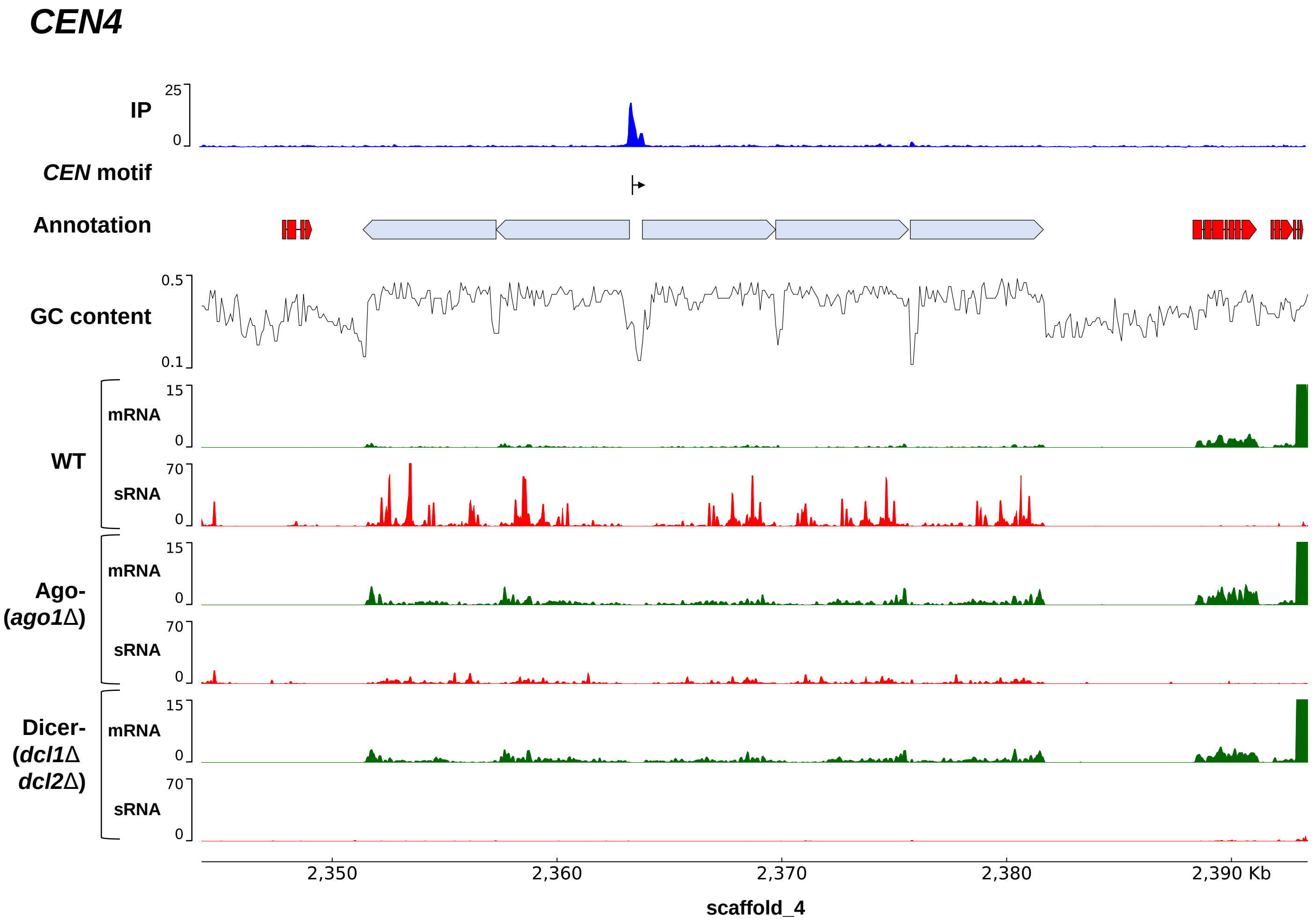
CEN4
IP
25
0
CEN motif
Annotation
GC content
0.5
0.1
WT
mRNA
15
0
sRNA
70
0
Ago-
(ago1Δ)
mRNA
15
0
sRNA
70
0
Dicer-
(dcl1Δ
dcl2Δ)
mRNA
15
0
sRNA
70
0
2,350
2,360
2,370
2,380
2,390 Kb
scaffold_4

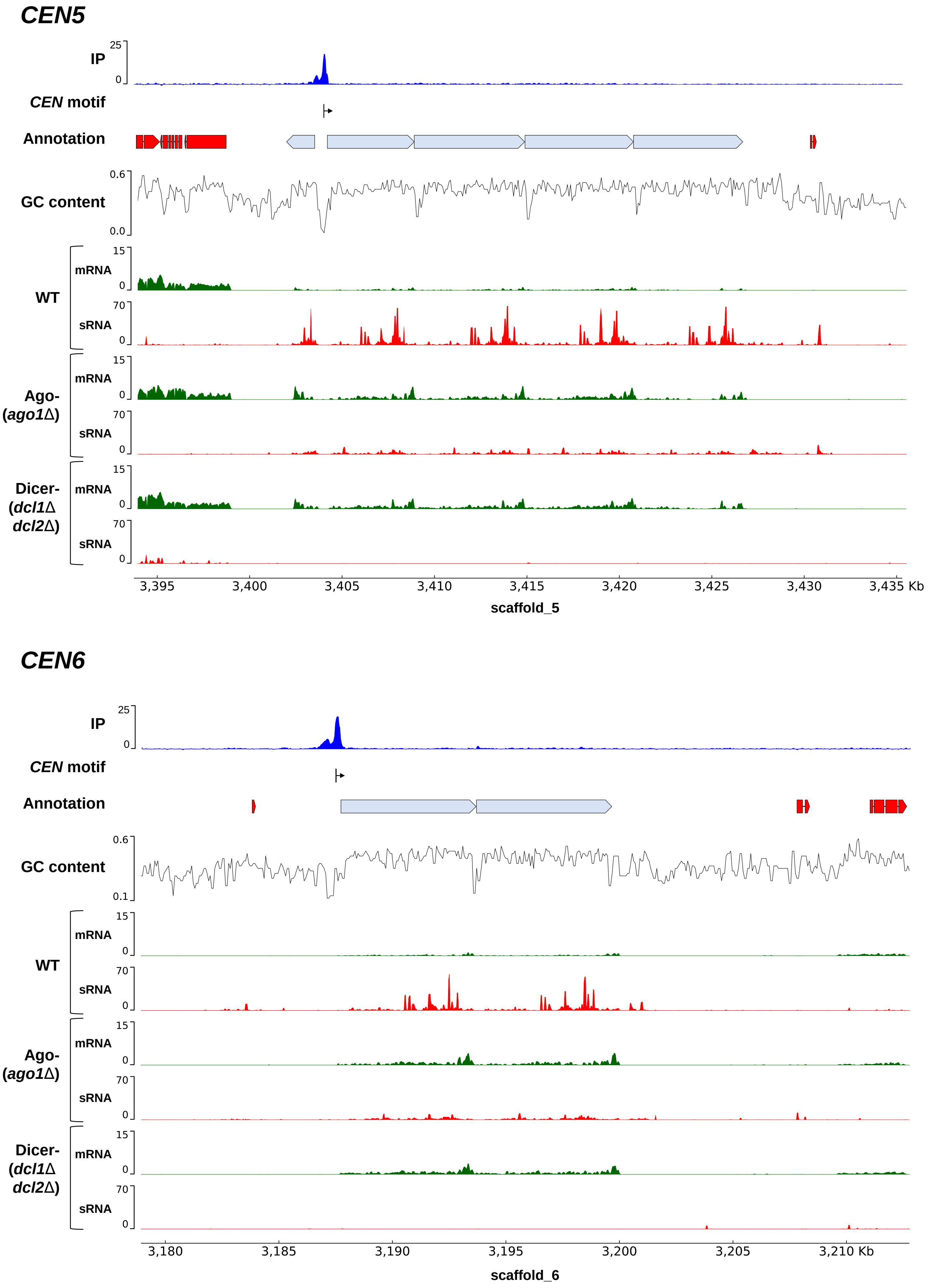
CEN5
IP
25
0
CEN motif
Annotation
GC content
0.6
0.0
WT
mRNA
15
0
sRNA
70
0
Ago-
(ago1Δ)
mRNA
15
0
sRNA
70
0
Dicer-
(dcl1Δ
dcl2Δ)
mRNA
15
0
sRNA
70
0
3,395
3,400
3,405
3,410
3,415
3,420
3,425
3,430
3,435 Kb
scaffold_5
CEN6
IP
25
0
CEN motif
Annotation
GC content
0.6
0.1
WT
mRNA
15
0
sRNA
70
0
Ago-
(ago1Δ)
mRNA
15
0
sRNA
70
0
Dicer-
(dcl1Δ
dcl2Δ)
mRNA
15
0
sRNA
70
0
3,180
3,185
3,190
3,195
3,200
3,205
3,210 Kb
scaffold_6

***CEN8***

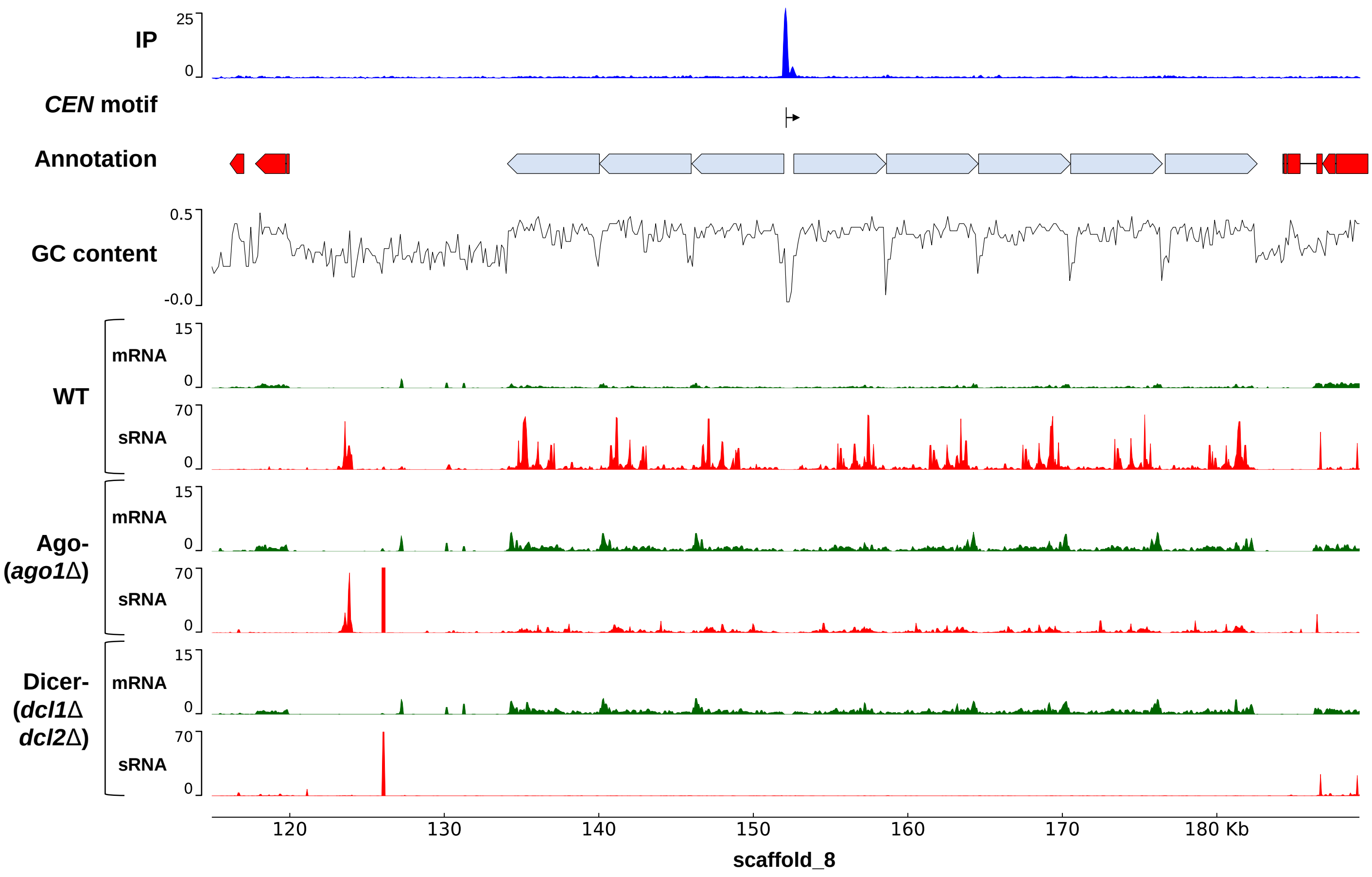
IP
25
0
*CEN* motif
Annotation
GC content
0.5
-0.0
WT
mRNA
15
0
sRNA
70
0
Ago-
(*ago1Δ*)
mRNA
15
0
sRNA
70
0
Dicer-
(*dcl1Δ*
*dcl2Δ*)
mRNA
15
0
sRNA
70
0
120
130
140
150
160
170
180 Kb
scaffold_8

***CEN9***

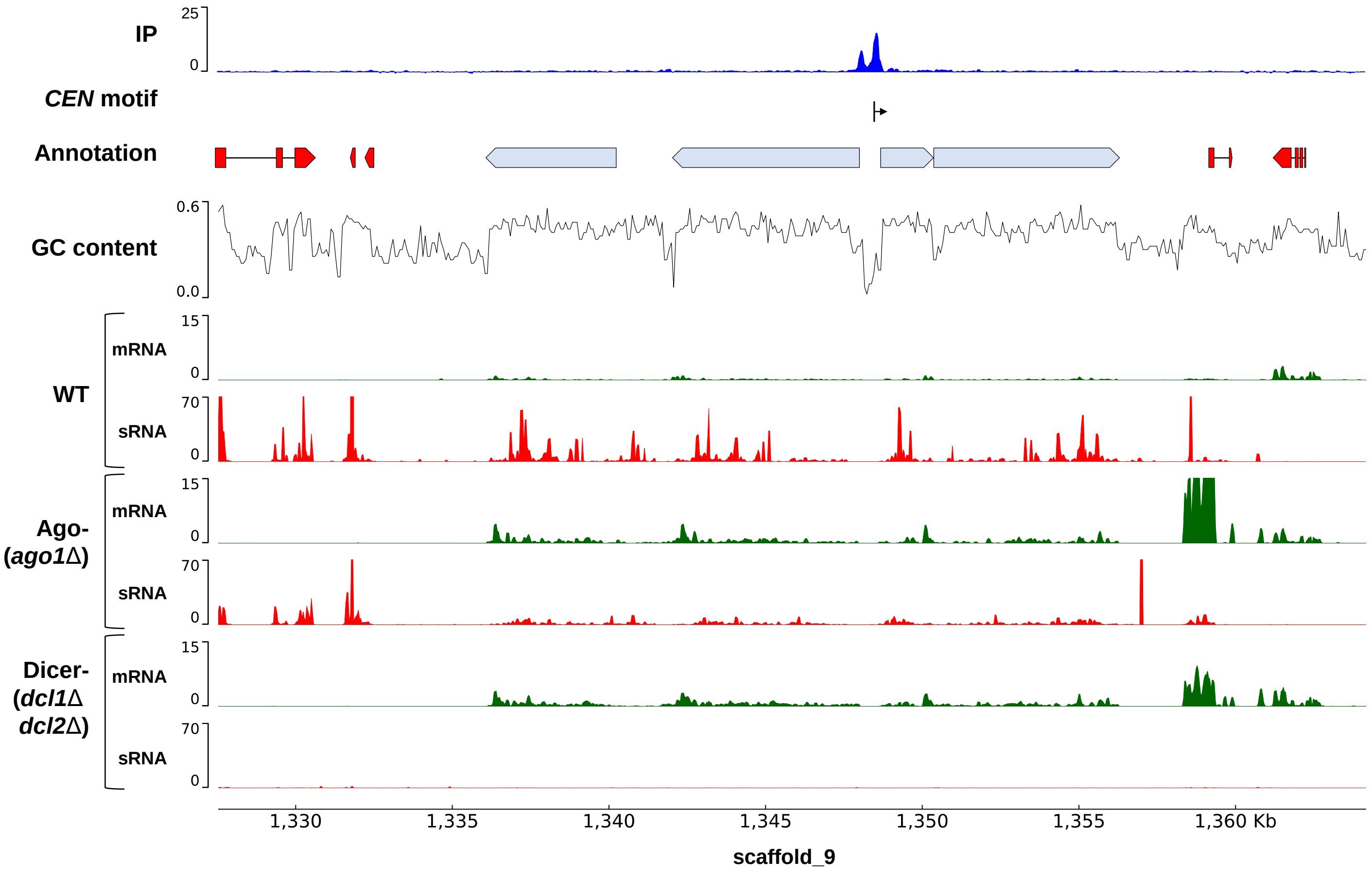
IP
25
0
*CEN* motif
Annotation
GC content
0.6
0.0
WT
mRNA
15
0
sRNA
70
0
Ago-
(*ago1Δ*)
mRNA
15
0
sRNA
70
0
Dicer-
(*dcl1Δ*
*dcl2Δ*)
mRNA
15
0
sRNA
70
0
1,330
1,335
1,340
1,345
1,350
1,355
1,360 Kb
scaffold_9

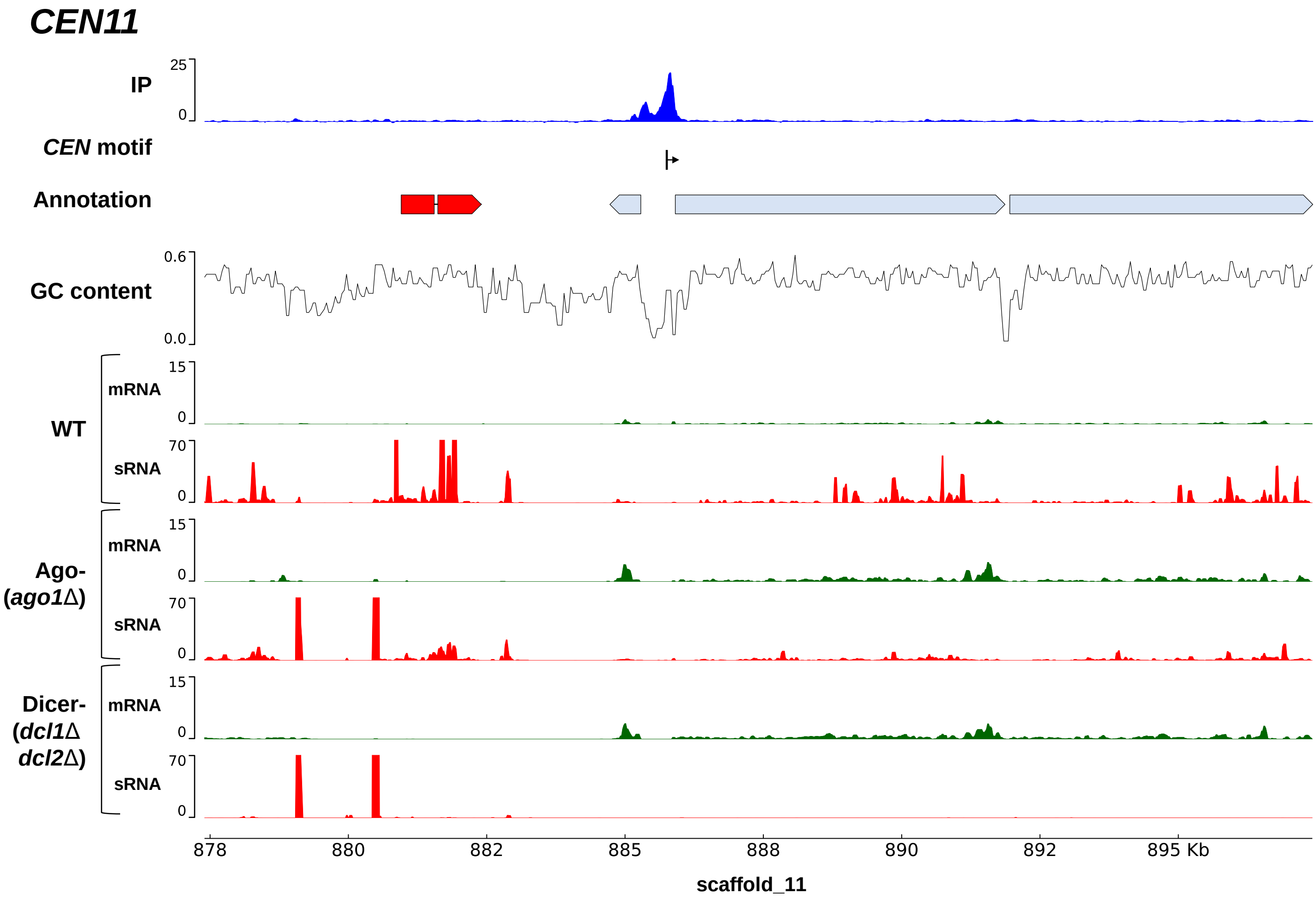

CEN11
IP
25
0
CEN motif
Annotation
GC content
0.6
0.0
WT
mRNA
15
0
sRNA
70
0
Ago-
(ago1Δ)
mRNA
15
0
sRNA
70
0
Dicer-
(dcl1Δ
dcl2Δ)
mRNA
15
0
sRNA
70
0
878
880
882
885
888
890
892
895 Kb
scaffold_11

### Supplementary Figure 7

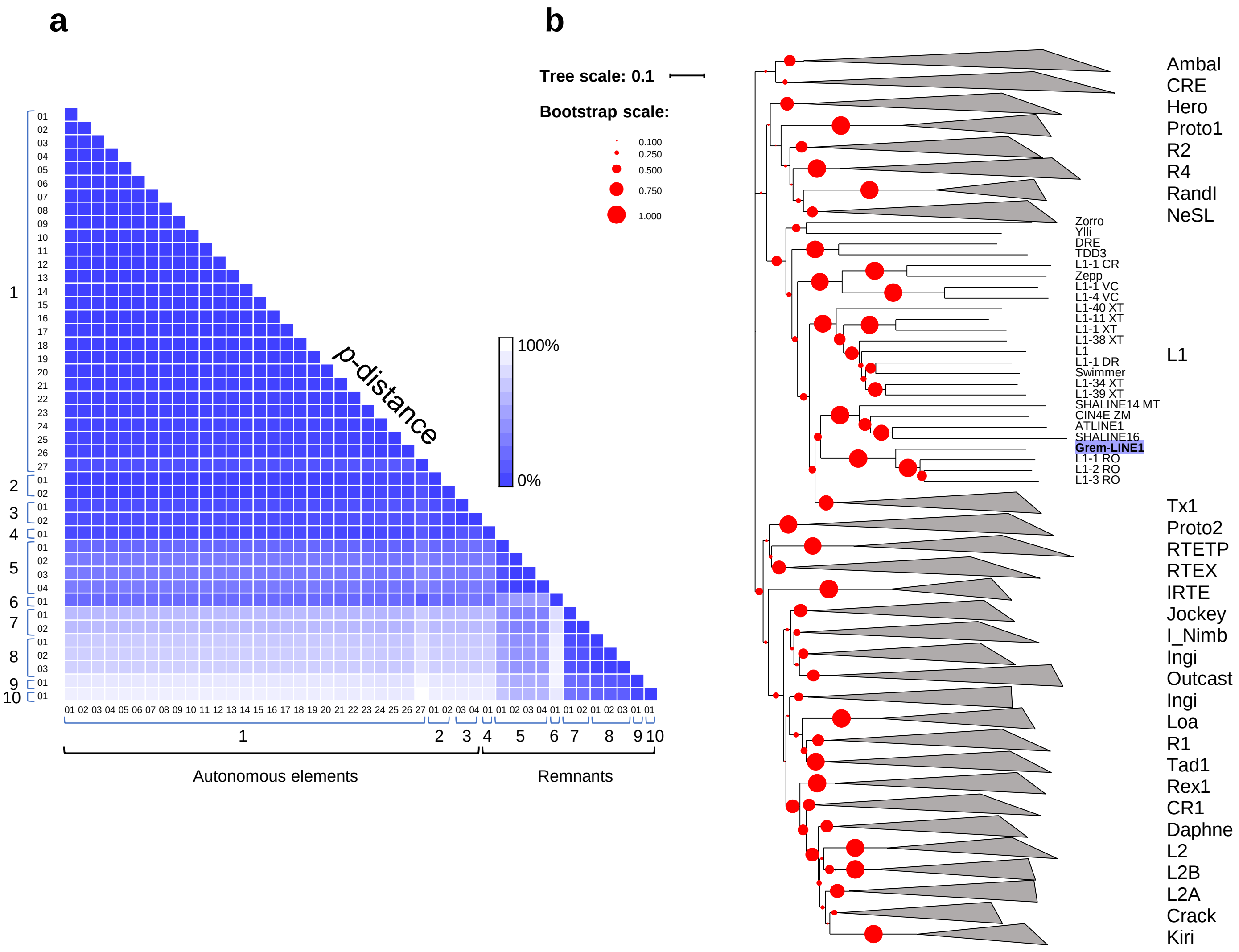
