## Supplementary Table 2 for "Early diverging fungus *Mucor circinelloides* lacks centromeric histone CENP-A and displays a mosaic of point and regional centromeres"

**Supplementary Table 2a. Primers used in the study**

| **Name** | **Sequence** | **Use** |
| --- | --- | --- |
| *mCherry*-F | ATGGTGAGCAAGGGCGAGGA | *mCherry* and *eGFP* amplification |
| *mCherry*-R | CTTGTACAGCTCGTCCATGC | *mCherry* and *eGFP* amplification |
| *mCherry*-R-*leuA* | gaatagagttggtagggagcaTTACTTGTACAGCTCGTCCATGC | *eGFP* tagging with marker *leuA* |
| *leuA*3kbFow | TGCTCCCTACCAACTCTATTC | *leuA* amplification |
| *leuA*3kbRev | GTCGAGTTGACCAGAATGTAC | *leuA* amplification |
| *pyrG*Fow2kb | TGCCTCAGCATTGGTACTTG | *pyrG* amplification |
| *pyrG*Rev2Kb | GTACACTGGCCATGCTATCG | *pyrG* amplification |
| P*zrt1*-F | cgatagcatggccagtgtacATCATCATCGATGTTTGTGCTGTC | Construction of pMAT1915 |
| P*zrt1*-R | CTCGAGATTTAGTTATTTTG | Construction of pMAT1915 |
| *carRP*-Inv-F | caagtaccaatgctgaggcaCCATATTGAGTCATCCTGCAACG | Construction of pMAT1915 |
| *carRP*-Inv-R | TACCACACATTGCAGACAGG | Construction of pMAT1915 |
| *hht4*-1 | tacatgggcccTCGCAATCATCCATGAAGTG | Hht4 tagging with eGFP at C-terminus |
| *hht4*-2 | tcctcgcccttgctcaccatAGAGCGTTCACCACGAAGA | Hht4 tagging with eGFP at C-terminus |
| *hht4*-3 | gtacattctggtcaactcgacATCATCATCTGATGTCTTTCTT | Hht4 tagging with eGFP at C-terminus |
| *hht4*-4 | tacatccgcggATGGCTCTGAAGTGATCCAC | Hht4 tagging with eGFP at C-terminus |
| *hhf1*-1 | tacatgtcgacACCAACAAACGTCACCTAGAGTAG | Hhf1 tagging with eGFP at C-terminus |
| *hhf1*-2 | tcctcgcccttgctcaccatTCCACCGAAACCGTAGAGGG | Hhf1 tagging with eGFP at C-terminus |
| *hhf1*-3 | gtacattctggtcaactcgacATCAAATCCCTCTGCTCATTCAC | Hhf1 tagging with eGFP at C-terminus |
| *hhf1*-4 | tacatctgcagCAACACGGGTGGTTTGGAG | Hhf1 tagging with eGFP at C-terminus |
| *hhf3*-1 | tacatgtcgacCCTAAAAGGGACAAAGATTATGGC | Hhf3 tagging with eGFP at C-terminus |
| *hhf3*-2 | tcctcgcccttgctcaccatTCCACCGAAACCGTAGAGGG | Hhf3 tagging with eGFP at C-terminus |
| *hhf3*-3 | gtacattctggtcaactcgacATGCAATTCATCATGCTTCTCAC | Hhf3 tagging with eGFP at C-terminus |
| *hhf3*-4 | tacatctgcagTGACAGGGCTTTCGCTTAGC | Hhf3 tagging with eGFP at C-terminus |
| *cnpT*-1 | cgaggtcgacggtatcgataCACGGCAGCAGCAGCAACATAC | CENP-T tagging with eGFP at C-terminus |
| *cnpT*-2 | tcctcgcccttgctcaccatTTCGTCTTCATTATCGTATCCACCG | CENP-T tagging with eGFP at C-terminus |
| *cnpT*-3 | gtacattctggtcaactcgacCTGTGATTGGTTGCCATGGTGG | CENP-T tagging with eGFP at C-terminus |
| *cnpT*-4 | caggaattcgatatcaagcTTGCTCGTGTATAGAACGAATCCAG | CENP-T tagging with eGFP at C-terminus |
| *mis12*-1 | tcctcgcccttgctcaccatTGGATCTGATGGCTGCTGG | Mis12 tagging with mCherry at C-terminus |
| *mis12*-2 | caaaataactaaatctcgagGATGCAAACCGACGAAAGCTA | Mis12 tagging with mCherry at C-terminus |
| *mis12*-3 | cctgtctgcaatgtgtggtaTAGTTTGAGAAGATTGTGGAGC | Mis12 tagging with mCherry at N-terminus |
| *mis12*-4 | ggcatggacgagctgtacaagATGCAAACCGACGAAAGCTA | Mis12 tagging with mCherry at N-terminus |
| *dsn1*-1 | tcctcgcccttgctcaccatTGGATCCTCCATCACAGAAGAT | Dsn1 tagging with mCherry at C-terminus |
| *dsn1*-2 | caaaataactaaatctcgagATGTCGGATAGACGCTTAAG | Dsn1 tagging with mCherry at C-terminus |
| *dsn1*-3 | cctgtctgcaatgtgtggtaCCCAACAGTAGAGCATCTTGG | Dsn1 tagging with mCherry at N-terminus |
| *dsn1*-4 | ggcatggacgagctgtacaagATGTCGGATAGACGCTTAAG | Dsn1 tagging with mCherry at N-terminus |
| *hht4*-ext-F | GCTACCTTGGATACCTGGAACA | *hht4* PCR confirmation |
| *hht4*-ext-R | CGAGTAAGGACGCCGTAGAC | *hht4* PCR confirmation |
| *hhf1*-Ext-F | GCTTCTTGACACCACCAGTAGAG | *hhf1* PCR confirmation |
| *hhf1*-Ext-R | GAATAGGTGGACAAGATGGGACT | *hhf1* PCR confirmation |
| *hhf3*-Ext-F | TATCTGTGAGGCTTCTTGACACC | *hhf3* PCR confirmation |
| *hhf3*-Ext-R | CGCTAAGTCCAAAGCAACTCTC | *hhf3* PCR confirmation |
| *cnpT*-Ext-F | CTTTTACCCTCTCAACCACGAG | *cnpT* PCR confirmation |
| *cnpT*-Ext-R | GCCTGTTTCAGATTGAGGGAAT | *cnpT* PCR confirmation |
| *carRP*-Ext-F | GGGCACATTGACGTAGAAGG | *mis12* and *dsn1* integration in the *carRP* locus PCR confirmation |
| *carRP*-Ext-R | GCTGTTGCTGTGCTAACATCAT | *mis12* and *dsn1* integration in the *carRP* locus PCR confirmation |

*Lowercase bases do not anneal to the the gene locus indicated in the table

**Underlined sequences are restriction sites

**Supplementary Table 2b. Primers used for H3 ChIP-qPCR**

| **Region of interest** | **Primer** | **Sequence (5’ to 3’)** | **Size of amplicon (bp)** | **Coordinates** |
| --- | --- | --- | --- | --- |
| *CEN2* core | F | GTTTCCTGAACGGGCTATTTG | 104 | 2: 4714295-4714315 |
|  | R | ACTGACAAAGTGTCCAAACCGA |  | 2:4714377-4714398 |
| *CEN2* 1L | F | GTACTGATGAAGCAAGAGGCG | 118 | 2: 4705002-4705022 |
|  | R | AGCTCTTTGTTCTCTGCTACCTTG |  | 2:4705096-4705119 |
| *CEN2* 2L | F | CCTTCCTGTTTTGATTGGCGG | 99 | 2:4707153-4707173 |
|  | R | TTGGCTTGCTAGAAGCACTTG |  | 2:4707231-4707251 |
| *CEN2* 1R | F | GCAAACGTTCATGGTAGTGCAAG | 123 | 2:4716962-4716984 |
|  | R | CTTCAGGTTGTGACAGATGTATGCA |  | 2:4717060-4717084 |
| *CEN2* 2R | F | TGGGAAATTCATCAAGGCCAGTGC | 98 | 2:4722407-4722430 |
|  | R | GAACCTCCTTTAGGGCCATGTTG |  | 2:4722482-4722504 |
| *CEN2* ORF-L | F | AAATTGCAGGACAGAAAAGACGC | 126 | 2:4692020-4692042 |
|  | R | CATGTCCAGCGCATCGCTTGATA |  | 2:4692123-4692145 |
| *CEN2* ORF-R | F | CAATTGACACGATGGGACTTTGAC | 102 | 2:4731524-4731547 |
|  | R | CAGTTTGACGCCGTATTGGAATG |  | 2:4731603-4731625 |
| *Far-CEN ORF* | F | CCTTGCCACTACCATCTGCTTC | 107 | 2:2231461-2231482 |
|  | R | ATCATCCATTCCCTCTTGTGCC |  | 2:2231546-2231564 |
