## Supplementary Table 3 for "Early diverging fungus *Mucor circinelloides* lacks centromeric histone CENP-A and displays a mosaic of point and regional centromeres"

**Supplementary Table 3. *M. circinelloides* f. *lusitanicus* strains used in this study**

| **Strain** | **Genotype** |
| --- | --- |
| MU402 | *leuA*^-^, *pyrG^-^* |
| MU636 | *leuA^-^*, *pyrG^+^* |
| MU837 | *hht4*-*eGFP*::*leuA, pyrG^+^* |
| MU838 | *hht4*-*eGPF*::*leuA, pyrG^+^* |
| MU839 | *hht4*-*eGPF*::*leuA, pyrG^+^* |
| MU840 | *carRP*::*mCherry*-*mis12*-*pyrG, leuA^-^* |
| MU841 | *carRP*::*mCherry*-*mis12*-*pyrG, leuA^-^* |
| MU842 | *carRP*::*mis12-mCherry*-*pyrG, leuA^-^* |
| MU843 | *carRP*::*mis12-mCherry*-*pyrG, leuA^-^* |
| MU844 | *carRP*::*mCherry*-*dsn1*-*pyrG, leuA^-^* |
| MU845 | *carRP*::*mCherry*-*dsn1*-*pyrG, leuA^-^* |
| MU846 | *carRP*::*dsn1-mCherry*-*pyrG, leuA^-^* |
| MU847 | *carRP*::*dsn1-mCherry*-*pyrG, leuA^-^* |
| MU851 | *carRP*::*mCherry*-*mis12*-*pyrG, hht4*-*eGFP*::*leuA* |
| MU852 | *carRP*::*mCherry*-*mis12*-*pyrG,* *hht4*-*eGFP*::*leuA* |
| MU853 | *carRP*::*mis12-mCherry*-*pyrG,* *hht4*-*eGFP*::*leuA* |
| MU854 | *carRP*::*mis12-mCherry*-*pyrG,* *hht4*-*eGFP*::*leuA* |
| MU855 | *carRP*::*mCherry*-*dsn1*-*pyrG, hht4*-*eGFP*::*leuA* |
| MU856 | *carRP*::*mCherry*-*dsn1*-*pyrG,* *hht4*-*eGFP*::*leuA* |
| MU857 | *carRP*::*dsn1-mCherry*-*pyrG,* *hht4*-*eGFP*::*leuA* |
| MU858 | *carRP*::*dsn1-mCherry*-*pyrG,* *hht4*-*eGFP*::*leuA* |
| MU657 | *cnpT*::*cnpT*-*eGFP*-*leuA,* *carRP*::*dsn1-mCherry*-*pyrG* |
| MU658 | *cnpT*::*cnpT*-*eGFP*-*leuA,* *carRP*::*dsn1-mCherry*-*pyrG* |
| MU659 | *cnpT*::*cnpT*-*eGFP*-*leuA,* *carRP*::*mis12-mCherry*-*pyrG* |
| MU660 | *cnpT*::*cnpT*-*eGFP*-*leuA,* *carRP*::*mis12-mCherry*-*pyrG* |
| MU661 | *hhf1*-*eGFP*::*leuA, pyrG^-^* |
| MU662 | *hhf3*-*eGFP*::*leuA, pyrG^-^* |
| MU663 | *hhf3*-*eGFP*::*leuA, pyrG^-^* |
| MU411 | *dcl1*::*pyrG,* *dcl2*::*leuA* |
| MU413 | *ago1*::*pyrG, leuA^-^* |
